# Standardized MNI-Guided TMS Yields Functional Similarity to Individualized T1-Guidance: Evidence from Behavioral, Anatomical, and Electromagnetic Levels

**DOI:** 10.64898/2026.05.14.725057

**Authors:** Hee-Dong Yoon, Hyeon-Ae Jeon

**Author notes:** Corresponding author. Seoul National University, Gwanak-ro 1, Gwanak-gu, Seoul, Korea 08826 E-mail address (Hyeon-Ae Jeon).

## Abstract

**Background:** Neuronavigation based on the standard MNI template (MNI-protocol) offers a cost-effective alternative to the gold-standard individualized T1-weighted MRI approach (T1-protocol). However, it remains unclear whether the reduced anatomical precision of the MNI-protocol compromises its functional efficacy, creating a critical need to verify protocol interchangeability.

**Objective:** We aimed to determine whether the MNI-protocol yields targeting efficacy comparable to the T1-protocol by specifically testing their functional and biophysical equivalence.

**Methods:** We employed a novel tri-level within-subject framework. The behavioral level assessed functional efficacy via the size congruity effect (SCE) during TMS to the right intraparietal sulcus (IPS). Anatomical accuracy (coil-to-cortex distances) and electromagnetic efficacy (E-field simulations) were evaluated across three distinct regions (right IPS, left dorsolateral prefrontal cortex, and left primary motor cortex) to assess regional generalizability.

**Results:** The MNI-protocol demonstrated functional similarity to the T1-protocol, yielding behavioral outcomes that were statistically indistinguishable. This functional equivalence was corroborated by electromagnetic analyses, which revealed nearly identical induced E-field magnitudes and spatial distributions across all three target regions. Although the T1-protocol achieved significantly shorter coil-to-cortex distances, this anatomical advantage did not confer any measurable functional benefit.

**Conclusion:** The MNI-protocol produced behavioral and electromagnetic outcomes equivalent to the T1-protocol. These findings validate the MNI-protocol as a scientifically sound and scalable alternative to individualized MRI-guided targeting, supporting its broader application in diverse research and clinical settings.

**Highlights:**

- Functional equivalence of MNI-vs. T1-guided TMS was systematically tested.
- A novel tri-level framework compared behavioral, anatomical, and E-field metrics.
- MNI- and T1-guided targeting yielded comparable behavioral and E-field outcomes.
- Anatomical proximity does not ensure better behavior or stronger E-field strength.
- MNI-guided targeting offers a robust, practical alternative to individual MRI.

## 1. Introduction

Accurate and consistent targeting is the cornerstone of effective transcranial magnetic stimulation (TMS); even minor coil deviations result in off-target stimulation, undermining both interpretability and reproducibility[1]. To ensure such precision, neuronavigation, utilizing frameless stereotaxy and real-time optical tracking, has become indispensable. For researchers, neuronavigation based on the standard MNI-ICBM152 template[2, 3] (MNI-protocol), offers a practical targeting solution. By using a population-averaged anatomical template rather than individual T1-weighted MRI scans, the MNI-protocol establishes a standardized framework for defining targets within a common coordinate system. This approach offers a distinct logistical advantage by bypassing separate MRI sessions, thus removing significant financial and operational barriers to research.

Conversely, when individualized precision is critical, the T1-protocol remains the gold standard, leveraging subject-specific T1 imaging for accurate neuronavigation[4, 5]. This approach affords high spatial fidelity and individual specificity[4, 5], enabling coil targeting guided by anatomical landmarks unique to each participant.

However, the T1-protocol imposes substantial logistical burdens: acquiring individual MRI scans demands significant time, resources, and coordination, often creating a bottleneck for large-scale research. In contrast, the MNI-protocol utilizing a standardized template circumvents these constraints, offering immediate accessibility and scalability. Yet, this pragmatic advantage comes at the cost of individualized anatomical precision[6], presenting a fundamental trade-off between logistical efficiency and spatial fidelity. This dichotomy necessitates empirical validation: if the MNI-protocol demonstrates functional comparability to the T1-protocol despite reduced spatial accuracy, the two methods may be considered interchangeable. Establishing such comparability would validate the MNI-protocol not merely as a convenient substitute, but as a scientifically sound framework combining practical scalability with functional efficacy comparable to the gold standard.

Definitive conclusions on protocol interchangeability remain elusive due to the fragmented nature of prior research. Previous studies used single-level analyses[4, 7–10] or separate cohorts[4, 9], precluding the direct integration required for a holistic conclusion. Therefore, a comprehensive assessment necessitates converging evidence across distinct levels from a unified, within-subject dataset. Furthermore, single-site evaluations fail to capture targeting variability driven by heterogeneous cortical depths[4, 7, 10, 11]. Since TMS efficacy relies on scalp-to-cortex distance[12], which varies substantially between superficial (e.g., the primary motor cortex[7, 10], M1) and deep (e.g., the dorsolateral prefrontal cortex[4], DLPFC) targets, findings from one locus cannot be reliably extrapolated to others. This leaves a critical gap in verifying protocol generalizability across diverse cortical topographies.

To bridge these gaps, we assessed the functional interchangeability of the T1- and MNI-protocols through a systematic, multi-level comparison. Using a within-subject design, we evaluated both protocols across three complementary analytical levels and three distinct cortical regions within the same participants. First, at the behavioral level, we quantified functional efficacy by measuring cognitive modulation during a number comparison task[8, 13, 14]. Second, at the anatomical level, we evaluated coil-to-cortex distances as a surrogate for spatial accuracy given its inverse relationship with magnetic field strength[15–17]. Third, at the electromagnetic level, we simulated induced electric-field (E-field) magnitude and propagation to capture biophysical variability driven by individual anatomy[18, 19]. Unlike prior single-region studies, our assessment spanned the right intraparietal sulcus (IPS), left DLPFC, and left M1, representing posterior, frontal, and mid-cortical locations. To our knowledge, this constitutes the first multifaceted, multi-region investigation designed to provide a comprehensive test of protocol interchangeability and standardized guidance for TMS research.

## 2. Methods

### 2.1. Experimental procedures

Figure 1A illustrates the overall experimental sessions. Participant information and the questionnaires used for screening[20, 21] are provided in Supplementary Materials 1 and 2, respectively. This study employed a single-blind, sham-controlled, within-subject design. While the first visit consisted exclusively of a T1-weighted MRI scan (T1 image), the remaining three visits involved separate TMS sessions: two active sessions (T1- and MNI-protocols) and one sham session (a control condition), with the order pseudo-randomized. See Supplementary Material 3 for details of MRI acquisition parameters. In each TMS session, participants performed a number comparison task (Fig. 1B)[8, 13, 14], identifying the numerically larger of two digits while ignoring physical size. Trials were congruent (numerical/physical size matched), incongruent (mismatched), or neutral (physically identical). This examined the size congruity effect (SCE), the typical response time (RT) advantage for congruent over incongruent trials[8, 14, 22]. RTs were measured from stimulus onset; SCE was quantified as the RT difference between incongruent and congruent conditions, indexing cognitive interference. See Supplementary Material 4 for task details.

**Fig. 1.**
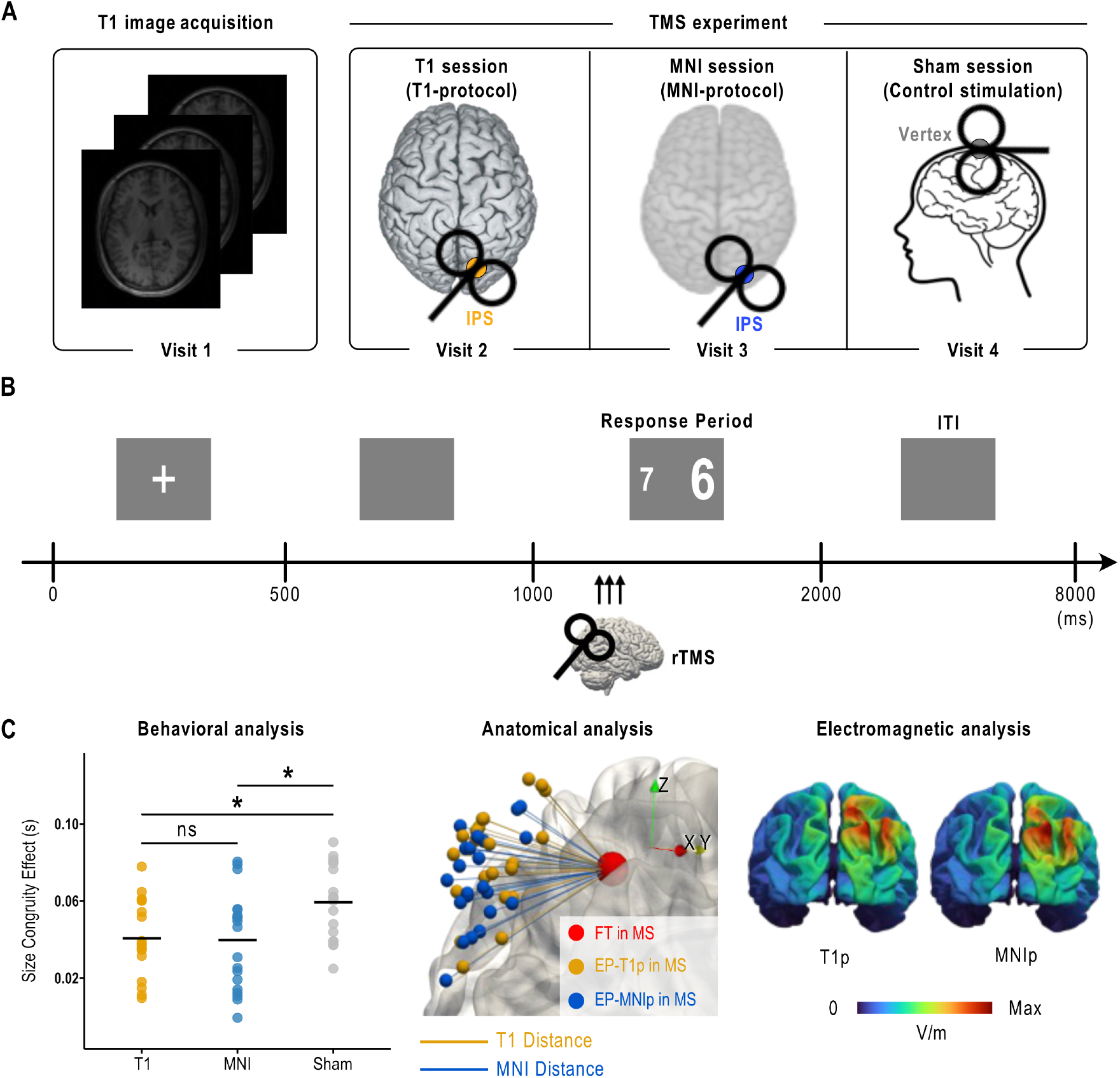
Experimental design and multi-level analytical framework. (A) Experimental sessions. Visit 1 involved T1-weighted MRI acquisition. Visit 2-4 comprised three TMS sessions: two active sessions and one sham control session. (B) Number comparison task. A digit pair was presented for 1000 ms, during which participants identified the numerically larger number. Three pulses of 10 Hz rTMS were delivered at 220, 320, and 420 ms post-stimulus onset. (C) Tri-level analysis. Behavioral: Functional efficacy was assessed via the SCE. Anatomical: Spatial accuracy was quantified as the Euclidean distance between the cortical functional target (FT) and the scalp entry point (EP). Electromagnetic: Biophysical efficacy was evaluated via E-field simulations to compare the magnitude and spatial distribution of the TMS-induced E-fields. Abbreviations: IPS, intraparietal sulcus; SCE, size congruity effect; MS, MNI space; T1p, T1-protocol; MNIp, MNI-protocol. ns, not significant; * *p* < 0.05.

Participants received event-related repetitive TMS (rTMS) over the right IPS, central to numerical magnitude processing[13, 14, 23, 24] and the SCE[8, 14, 22]. The target (MNI: x = 25.63, y = −66.73, z = 45.28) was defined based on peak SCE activations in previous studies[14, 25]. Triple-pulse 10 Hz rTMS was delivered at 220, 320, and 420 ms post-stimulus to disrupt size-number interference[22]. Targeting the right IPS during this window aimed to perturb size-number interference. This paradigm was used in both active sessions (T1 and MNI), whereas sham stimulation targeted the vertex. TMS effectiveness was indexed by the SCE difference between active and sham sessions. See Supplementary Material 5 for TMS stimulation parameters.

### 2.2. Neuronavigation and targeting protocols

The fundamental distinction between the MNI- and T1-protocols lies in spatial referencing (Fig. 2A). In the MNI-protocol, the MNI template was registered and individualized to the participant by scanning predefined anatomical landmarks (e.g., nasion, preauricular points) and the head surface with a digitizing pointer. This procedure leverages linear transformations and non-linear deformations to adapt the population-averaged template to the participant’s specific head geometry. Consequently, MNI space (MS) functioned as a unified environment for both target definition and the real-time visualization of coil poses (position and orientation) during navigation.

**Fig. 2.**
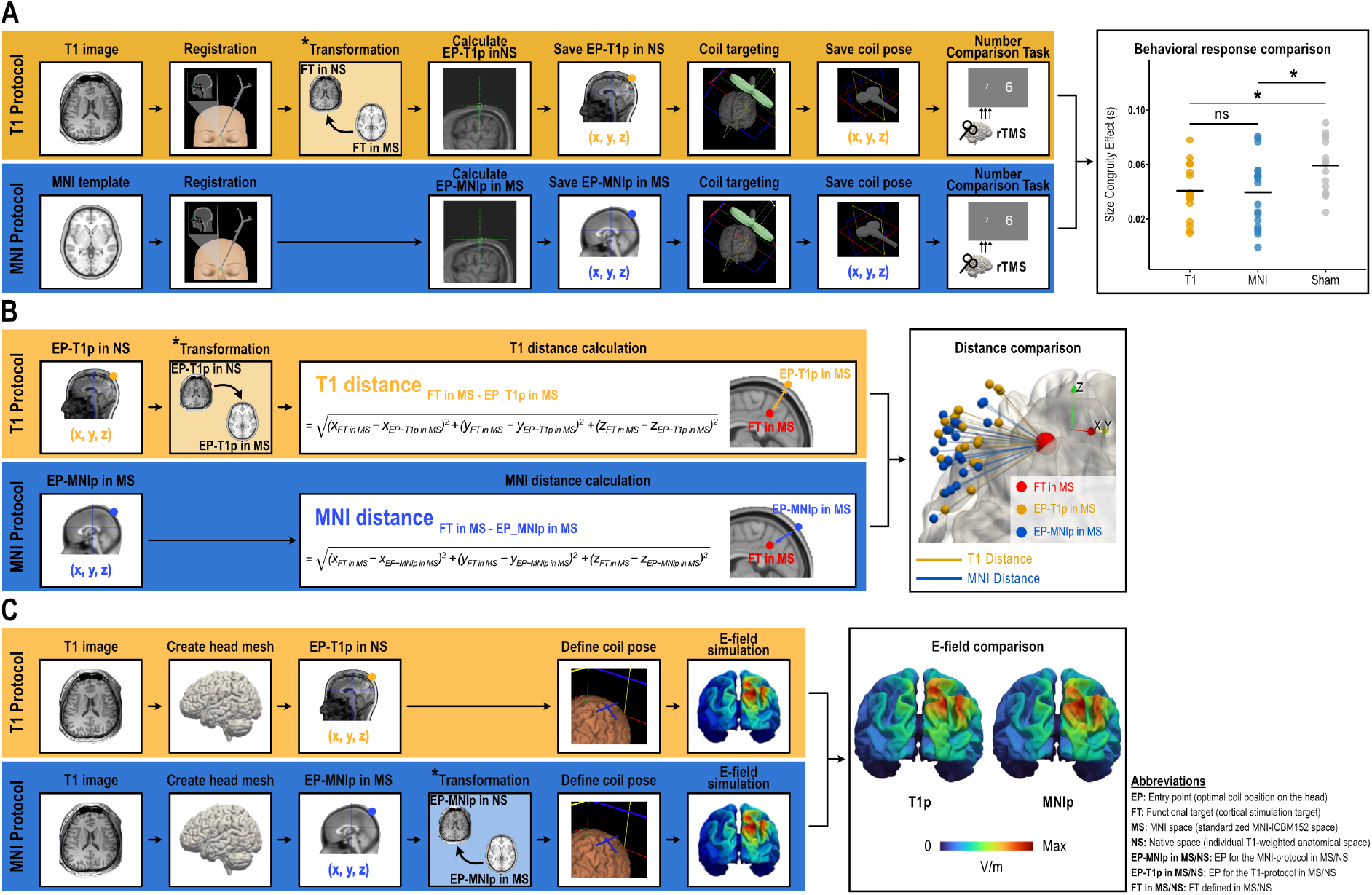
Methodological workflow for the multi-level comparison of T1- and MNI-protocols. The schematic illustrates the T1-protocol (yellow panels) and MNI-protocol (blue panels) across three distinct levels of analysis. Asterisks (*) denote the coordinate transformation steps. (A) Behavioral level. Participants received rTMS guided by either their individual T1 image or the MNI template. In the T1-protocol, FT coordinates, originally defined in MS, were inverse-normalized to the participant’s NS (FT in MS → FT in NS) to calculate the individualized EP in NS. In contrast, the MNI-protocol required no transformation because FT and EP were defined directly within the standardized MS. Following targeting, behavioral efficacy was assessed via the size congruity effect during the number comparison task. (B) Anatomical level. To ensure a standardized comparison, coil-to-cortex distances were quantified in the MS. For the T1-protocol, the EP calculated in NS (EP-T1p in NS) was transformed into MS (EP-T1p in NS → EP-T1p in MS), whereas for the MNI-protocol, EP was defined in MS (EP-MNIp in MS). Euclidean distances between the FT and the protocol-specific EPs were then computed in MS. (C) Electromagnetic level. Simulations were performed in NS, using head meshes generated from individual T1 images. For the T1-protocol, coil placement was guided by the EP-T1p defined in NS. For the MNI-protocol, the EP-MNIp defined in MS was transformed into NS (EP-MNIp in MS → EP-MNIp in NS) to replicate the physical stimulation geometry on the individual anatomy.

In the T1-protocol, each participant’s T1 image served as the navigation space, allowing coil positions to be visualized directly on the subject-specific anatomy. Accordingly, right IPS target coordinates, defined in MS, were transformed into each participant’s native space (NS) for individualized targeting. The transformation methodology, its validation procedure, and the results confirming its robustness are detailed in Supplementary Materials 6, 7, and 8–10, respectively.

Target coordinates for both protocols were imported into the Localite neuronavigation system (Localite TMS Navigator, Localite GmbH, Sankt Augustin, Germany) to calculate the TMS entry point (EP). Defined as the scalp position providing the shortest path to the cortical target[26], the EP served as the anchor for subsequent anatomical (coil-to-cortex distance) and electromagnetic (E-field) analyses. See Supplementary Material 11 for EP calculation.

### 2.3. Analysis

We investigated the functional interchangeability of the T1- and MNI-protocols for TMS targeting, using a systematic tri-level analytical framework (Fig. 1C). This design was engineered to capture three dimensions of targeting performance: functional efficacy via cognitive tasks (SCE), spatial accuracy via anatomical distances (coil-to-cortex distance), and biophysical induction via E-field simulations. By integrating these multi-level indices, we evaluated convergence of findings across levels to robustly validate the biophysical and functional equivalence of the two protocols.

#### 2.3.1. The behavioral level: Size congruity effect (SCE)

We evaluated whether the T1- and MNI-protocols differently modulated cognitive interference during numerical processing, indexed by the SCE. Session-wise SCE values were analyzed via a linear mixed-effects model [*SCE ~ SESSION + (1*|*participant)*], using Kenward-Roger method for degrees of freedom and Tukey-adjusted pairwise contrasts. To isolate condition-specific interference from global shifts in response speed, RTs were assessed using a two-way repeated measures ANOVA with within-subject factors SESSION (T1-protocol, MNI-protocol, sham) and CONDITION (congruent, incongruent, neutral). Behavioral data preprocessing procedures are detailed in Supplementary Material 12.

#### 2.3.2. The anatomical level: Coil-to-cortex distance

To assess spatial accuracy, we quantified the Euclidean distance between the scalp EP and its corresponding functional target (FT), that is, the coil-to-cortex distance (Fig. 2B). As the EP represents the optimal scalp position for coil placement, a minimized EP-to-FT distance reflects superior targeting accuracy, ensuring the coil is in closer physical proximity to the intended cortical target. To control for individual anatomical variability and head shape differences, all distances were computed in standardized MS. Specifically, the T1-protocol distance used an EP originally determined in the participant’s NS and then spatially transformed into MS, while the MNI-protocol distance utilized an EP defined directly within MS.

To ensure regional generalizability, these distances were computed for the right IPS, left DLPFC, and left M1 using standardized MNI coordinates from previous studies (right IPS: x = 25.63, y = −66.73, z = 45.28; left DLPFC: x = −30, y = 43, z = 23; left M1: x = −37, y = −21, z = 58)[27–30]. Within each region, T1 and MNI distances were compared using paired-sample t-tests. The full computation pipeline is detailed in Supplementary Material 13.

#### 2.3.3. The electromagnetic level: E-field simulation

##### Simulation pipeline and biophysical validity

To assess T1- and MNI-protocol comparability in electromagnetic effects, we simulated TMS-induced E-fields using SimNIBS (v3.2.1; Fig. 2C). Simulations were conducted across three cortical regions (right IPS, left DLPFC, and left M1) to ensure the regional generalizability. Critically, all simulations were performed in each participant’s NS to ensure biophysical validity by preserving subject-specific anatomical features, such as cortical morphology, which fundamentally influence E-field strength and distribution[31, 32]. Accordingly, for the T1-protocol, simulations utilized EPs calculated directly in NS. For the MNI-protocol, EPs were inverse-normalized from MS into each individual’s NS to enable valid comparison within the same biophysical environment. Detailed simulation workflows and individually constructed head models are provided in Supplementary Materials 14 and 15, respectively.

##### “Real” vs. “Virtual” TMS framework

We implemented a dual approach, “Real” and “Virtual” TMS, to overcome data limitations while ensuring targeting accuracy. “Real TMS” denotes simulations driven by coil poses empirically recorded during the actual TMS sessions (available only for the right IPS), providing a high-fidelity reconstruction of the stimulation delivered to participants. Conversely, “Virtual TMS” utilizes coil poses manually configured via the neuronavigation interface on subject-specific head models. This virtual approach was necessary for the left DLPFC and left M1, where recorded coil poses were unavailable. To validate this method, we demonstrated at the right IPS that manually defined coil poses (Virtual) successfully reproduced E-field estimates derived from empirical simulations (Real). Consequently, to maintain methodological consistency across all regions, Virtual TMS was adopted for the primary cross-region analysis. Approach-specific coil pose operationalization and validation procedures are detailed in Supplementary Material 16.

##### Stepwise comparative analysis

1. **Protocol similarity at the target (Real T1 vs. Real MNI):** We first compared E-field simulations using coil poses recorded at the right IPS during the behavioral TMS sessions to assess protocol similarity in the induced fields.
2. **Validation of Virtual TMS (Real TMS vs. Virtual TMS):** We compared empirically driven (Real) and manually defined (Virtual) simulations at the right IPS to confirm that the virtual approach accurately matches empirical E-field patterns.
3. **Generalizability of E-field equivalence (Virtual T1 vs. Virtual MNI):** Leveraging the validated virtual approach, we compared virtual E-field simulations across all three regions (right IPS, left DLPFC, and left M1) to determine whether protocol similarity in the induced fields generalizes across distinct anatomical locations.

##### Outcome metrics

For each target and participant, E-field magnitudes were quantified using two complementary metrics. First, we extracted the mean E-field magnitude within a region of interest (ROI), operationalized as a 5 mm-radius sphere centered on the FT coordinates using SimNIBS[33] (Fig. 3A; see Supplementary Material 17), and compared it across protocols using paired-sample t-tests. Second, to assess the spatial extent, we computed the total cortical surface area (in mm^2^) where the E-field exceeded 50% of the group-averaged maximum magnitude (see Supplementary Material 18).

**Fig. 3.**
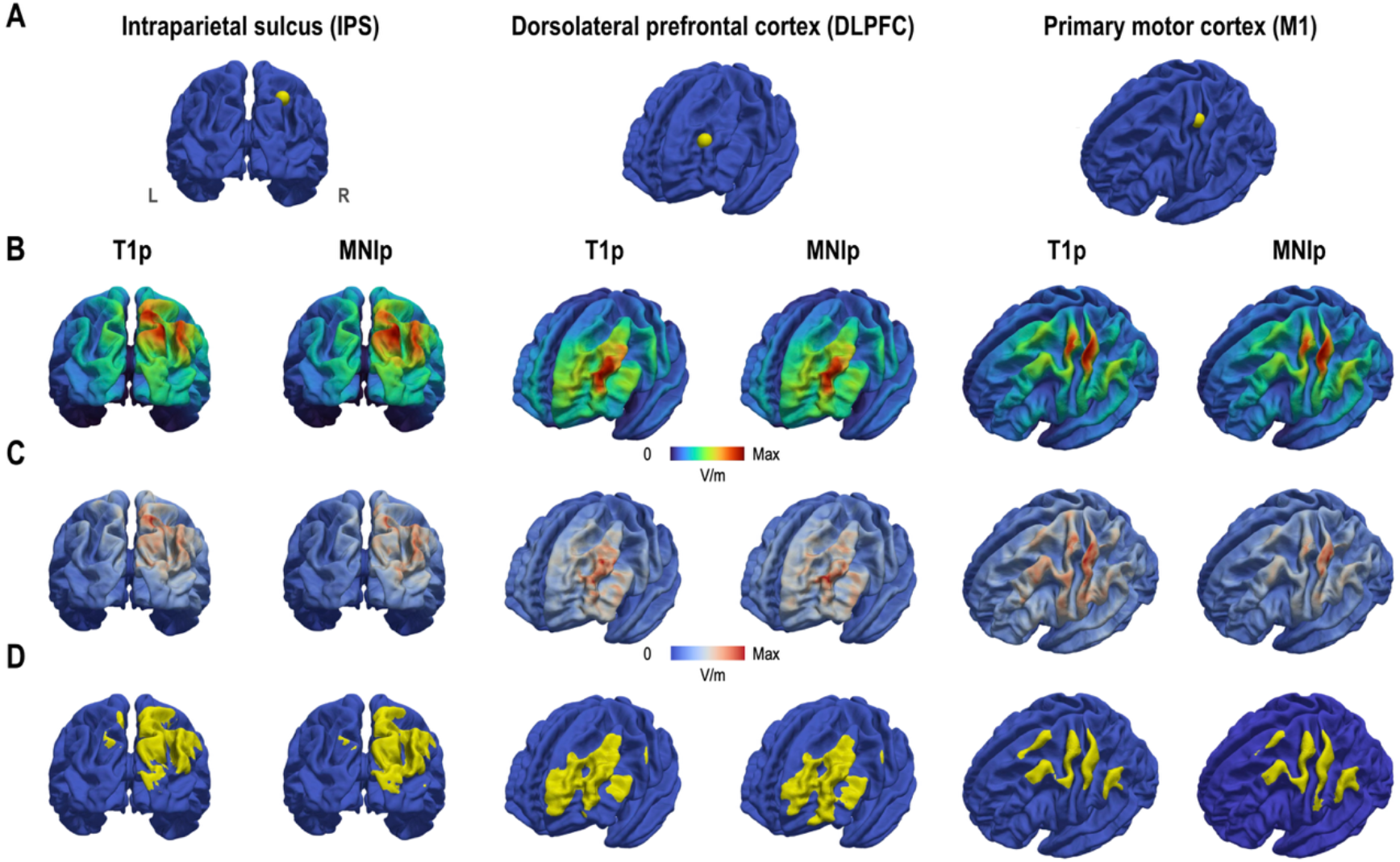
Comparative E-field modeling across three cortical regions: T1-versus MNI-protocols (Virtual TMS). (A) ROI definition. Yellow spheres indicate the 5-mm radius ROIs. (B) Group-averaged E-field magnitude. Spatial distribution of the mean E-field (V/m) simulated under the T1-protocol (T1p) and MNI-protocol (MNIp), projected onto the *fsaverage* surface. The colormap is scaled from 0 V/m to protocol-specific maximums (IPS: T1p 52, MNIp 53; DLPFC: T1p 71, MNIp 73; M1: T1p 71, MNIp 69). (C) Inter-subject variability. Variability is expressed as standard deviation (SD; V/m), scaled to protocol-specific maximums (IPS: T1p 24, MNIp 26; DLPFC: T1p 26, MNIp 28; M1: T1p 21, MNIp 28). (D) Visualization of E-field spread. Yellow areas denote the spatial extent where the group-averaged E-field exceeds 50% of the peak magnitude.

## 3. Results

### 3.1. Behavioral level: Functional equivalence of T1- and MNI-protocols

We analyzed size congruity effect (SCE) scores across all TMS sessions. A linear mixed-effects model revealed a significant main effect of SESSION (*F*(2, 55) = 5.20, *p* = 0.009; Fig. 4A). Critically, estimated pairwise comparisons showed no significant difference in SCE modulation between T1- and MNI-protocols (Δ = 0.001s, *p* = 0.990), underscoring their functional similarity. Notably, both active protocols produced a significant reduction in the SCE compared to the sham session (T1 vs. sham: Δ = −0.019s, *p* = 0.027; MNI vs. sham: Δ = −0.020s, *p* = 0.018). These findings confirm that active TMS over the right IPS reliably modulated numerical magnitude processing, and more importantly, that the T1- and MNI-protocols were functionally indistinguishable at the behavioral level.

**Fig. 4.**
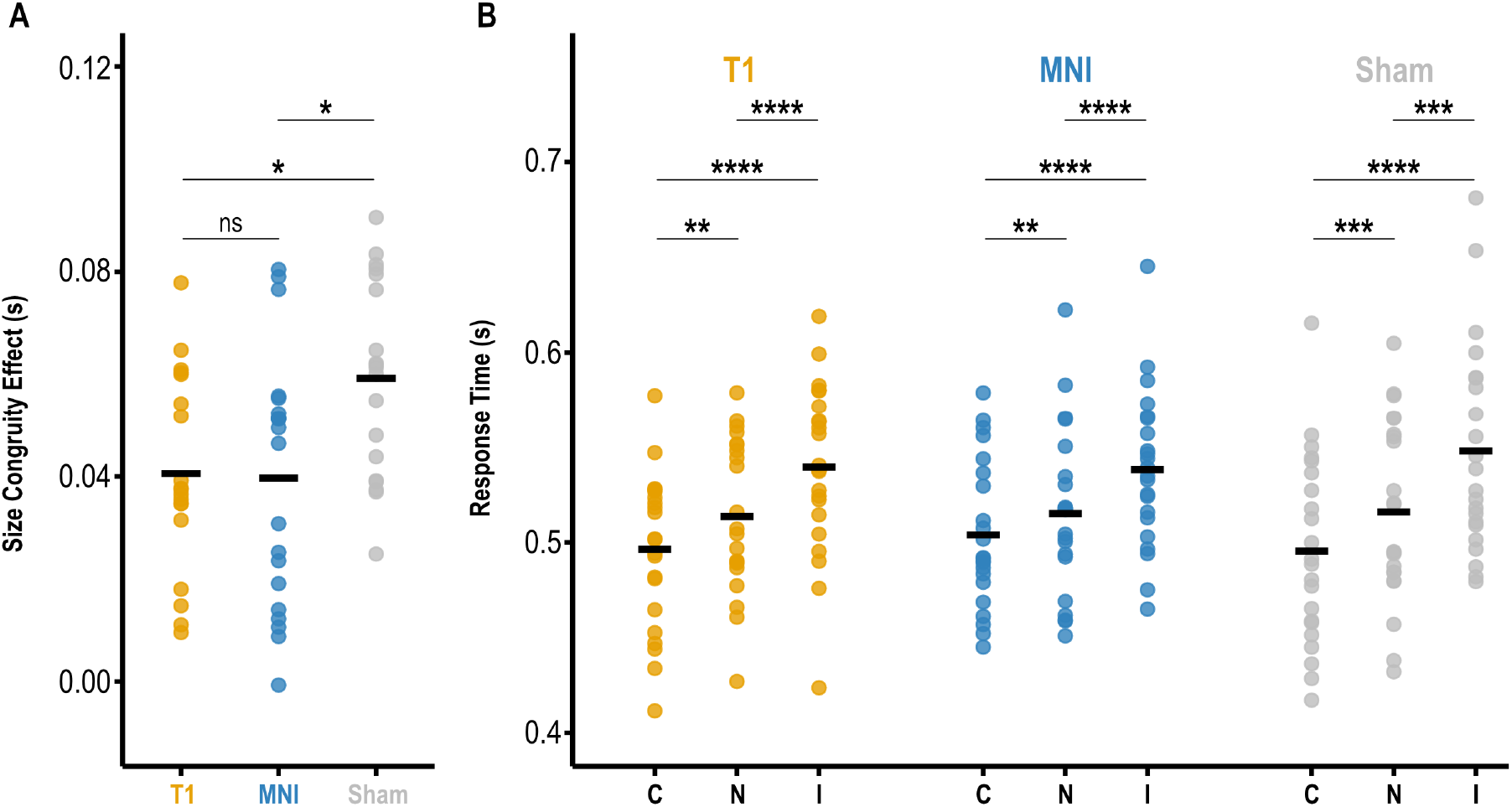
Behavioral comparison of T1- and MNI-protocols. (A) Functional equivalence in size congruity effect. Individual (dots) and group-mean (horizontal bars) SCE values are shown for the T1 (yellow), MNI (blue), and sham (gray) sessions. (B) Response times by condition. Mean RTs are stratified by trial condition (C: congruent, N: neutral, I: incongruent). ns, not significant; * *p* < 0.05; ** *p* < 0.01; *** *p* < 0.001; **** *p* < 0.0001 (Bonferroni-corrected).

Complementary analysis of RT data using a two-way repeated-measures ANOVA confirmed a robust main effect of CONDITION (*F*(2, 42) = 139.07, *p* < 0.001; Fig. 4B), validating the stable presence of the SCE across all sessions. Participants responded significantly faster in congruent trials (*t*_(21)_ = –7.07, *p* < 0.001) and slower in incongruent trials (*t*_(21)_ = 8.67, *p* < 0.001) relative to neutral trials. Furthermore, the absence of a significant main effect of SESSION (*F*(2, 42) = 0.10, *p* = 0.90) or SESSION × CONDITION interaction (*F*(4, 84) = 1.34, *p* = 0.26) indicates that the specific targeting protocol did not induce global shifts in response speed or alter the fundamental RT patterns of the task.

### 3.2. Anatomical level: Superior spatial accuracy of the T1-protocol vs. practicality of the MNI-protocol

Across all three cortical regions, the MNI-protocol yielded significantly greater coil-to-cortex distances than the T1-protocol: right IPS (*t*_(21)_ = 6.13, *p* < 0.001), left DLPFC (*t*_(21)_ = 7.49, *p* < 0.001), and left M1 (*t*_(21)_ = 4.53, *p* < 0.001). Summary statistics and individual data values indicating this robust pattern are provided in Table 1 and Supplementary Material 19, respectively. The T1-protocol achieves superior spatial fidelity by utilizing individual brain images that preserve subject-specific cortical geometry. The MNI-protocol’s reliance on a population-averaged template produces systematically larger coil-to-cortex distances due to its inability to account for individual anatomical characteristics. Despite the significant protocol-wise differences in physical proximity to the target, the behavioral outcomes remained functionally comparable.

**Table 1.**
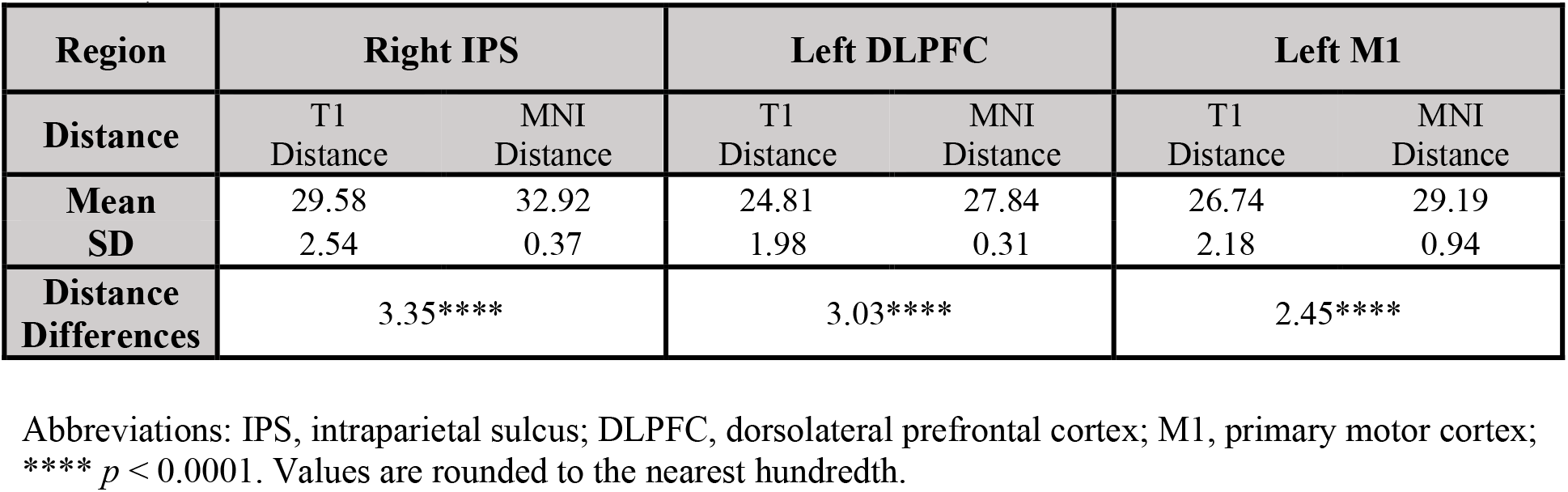
T1 distance and MNI distance (mm).

| Region | Right IPS |  | Left DLPFC |  | Left M1 |  |
| --- | --- | --- | --- | --- | --- | --- |
| Distance | T1 Distance | MNI Distance | T1 Distance | MNI Distance | T1 Distance | MNI Distance |
| Mean | 29.58 | 32.92 | 24.81 | 27.84 | 26.74 | 29.19 |
| SD | 2.54 | 0.37 | 1.98 | 0.31 | 2.18 | 0.94 |
| Distance Differences | 3.35**** |  | 3.03**** |  | 2.45**** |  |

Abbreviations: IPS, intraparietal sulcus; DLPFC, dorsolateral prefrontal cortex; M1, primary motor cortex; \*\*\*\* $p < 0.0001$ . Values are rounded to the nearest hundredth.

### 3.3. Electromagnetic level: E-field simulations

#### 3.3.1. Biophysical equivalence of T1- and MNI-protocol E-fields

At the primary experimental target (right IPS), simulated E-field magnitudes based on empirically recorded coil poses demonstrated striking consistency across protocols. The mean E-field magnitude was 29.04 ± 9.14 V/m for the T1-protocol and 28.91 ± 6.73 V/m for the MNI-protocol, with no statistically significant difference (*t*_(21)_ = 0.10, *p* = 0.92). Participant-wise E-field distribution maps and mean magnitudes are provided in Supplementary Materials 20 and 21, respectively. Despite differences in physical coil-to-cortex proximity, both protocols delivered equivalent electromagnetic magnitude to the target, challenging the assumption that closer targeting guarantees superior functional or physiological efficacy in TMS.

#### 3.3.2. Validation of Virtual TMS

We validated the robustness of the Virtual TMS approach at the right IPS. A comparison between Real TMS (driven by recorded coil poses) and Virtual TMS (utilizing manually configured poses) showed that participant-wise E-field magnitudes were significantly indistinguishable (*t*_(21)_ = – 0.96, *p* = 0.35; Supplementary Material 22). As illustrated in Figure 5, both modeling approaches yielded highly convergent E-field spatial patterns. Consequently, E-field estimates derived from manually configured virtual coil poses provide a reliable proxy for empirical stimulation, justifying the use of Virtual TMS for evaluating regional generalizability in subsequent analyses.

**Fig. 5.**
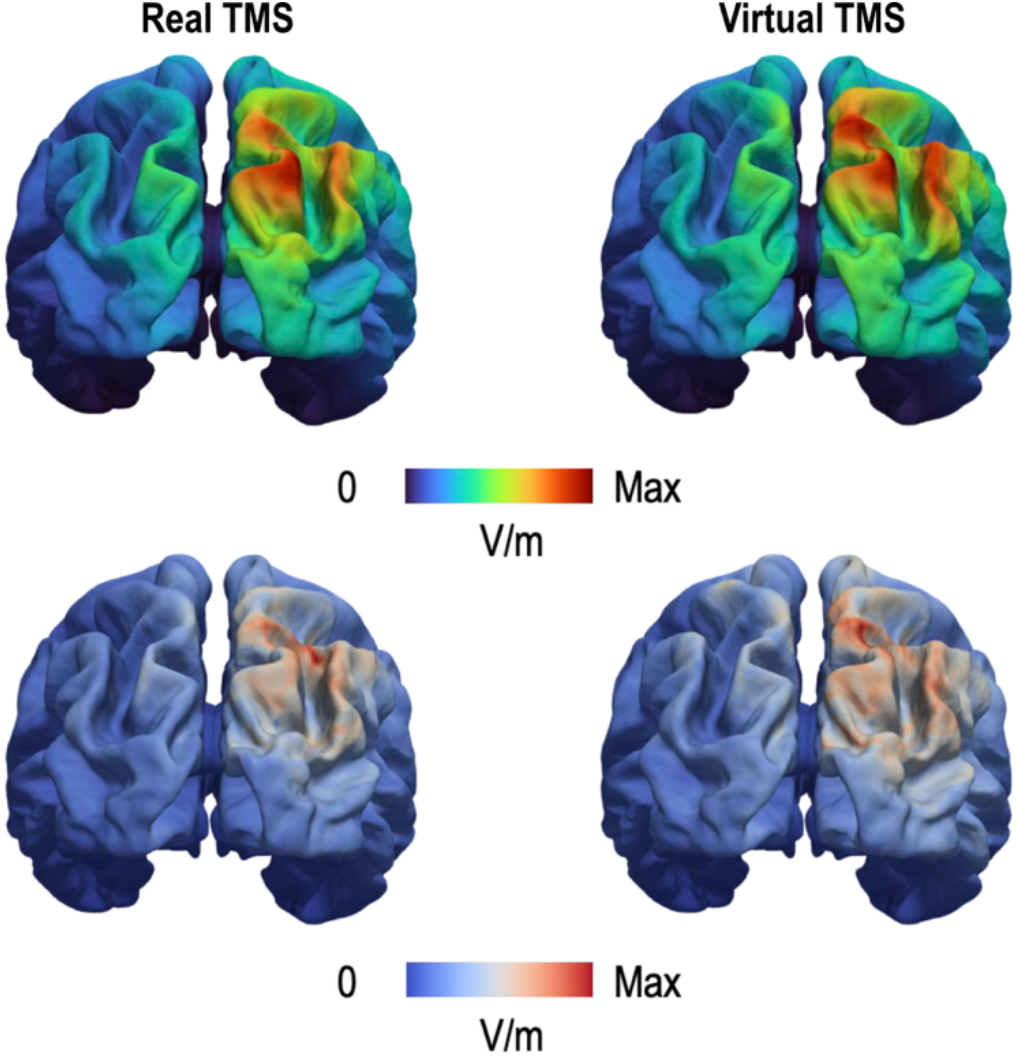
Methodological validation of E-field simulations: Real versus Virtual TMS at the right IPS. Top row: Group-averaged E-field magnitude (V/m) overlaid on the *fsaverage* brain surface for real TMS (left) and virtual TMS (right). The colormap is scaled from 0 V/m to the approach-specific maximum (real TMS: 52; virtual TMS: 52). Bottom row: Inter-subject variability maps expressed as standard deviation (SD; V/m). The colormap is scaled from 0 V/m to the approach-specific maximum (real: 28; virtual: 24). Abbreviations: IPS, intraparietal sulcus.

#### 3.3.3. Regional generalizability: Interchangeability of protocols across cortical sites

We evaluated regional generalizability across three distinct cortical areas: the right IPS, left DLPFC, and left M1 (Fig. 3A). Mean E-field magnitudes were excracted within subject-specific ROIs to determine whether the two protocols delivered comparable physiological dose (Fig. 3B-C). The induced E-field magnitudes were closely matched and did not differ significantly between protocols in any region: right IPS (30.18 ± 6.51 V/m vs. 29.06 ± 7.49 V/m; *t*_(21)_ = 0.69, *p* = 0.50), left DLPFC (39.71 ± 8.39 V/m vs. 39.32 ± 9.99 V/m; *t*_(21)_ = 0.21, *p* = 0.84), and left M1 (30.77 ± 5.84 V/m vs. 30.20 ± 8.01 V/m; *t*_(21)_ = 0.33, *p* = 0.75).

Beyond absolute magnitude, we characterized the spatial extent of E-field propagation by quantifying the total cortical surface area where E-field exceeded 50% of the group-averaged maximum magnitude (Fig. 3D). This spatial profile remained highly comparable between the T1- and MNI-protocols across all targets: IPS (4405 mm^2^ vs. 4613 mm^2^), DLPFC (3316 mm^2^ vs. 3353 mm^2^), and M1 (2349 mm^2^ vs. 2556 mm^2^). Thus, the two protocols produce equivalent electromagnetic effects regardless of the specific anatomical location or target depth.

## 4. Discussion

This study establishes the functional equivalence of individualized (T1) and standardized (MNI) TMS targeting. Utilizing a tri-level framework, we demonstrate that despite the T1-protocol’s superior spatial precision, the MNI-protocol yields comparable behavioral and electromagnetic efficacy. To our knowledge, this is the first multi-level validation of protocol interchangeability, confirming MNI-guidance as a scientifically sound, scalable alternative to individualized MRI.

### 4.1. Functional similarity of T1- and MNI-protocols

Our results demonstrate robust functional similarity between the T1- and MNI-protocols, converging across behavioral and electromagnetic levels. TMS-induced behavioral modulation (SCE; Fig. 4A) was statistically indistinguishable between protocols, and electromagnetic metrics (E-field; Fig. 3) were matched across all tested regions (right IPS, left DLPFC, and left M1). By leveraging physiologically grounded E-field modeling based on Faraday’s law of induction[12, 34], we provide a standardized, quantitative validation that transcends behavioral observation alone. This convergence is particularly noteworthy, as it aligns the protocol’s direct physical cause (induced E-field) with its ultimate functional outcome (behavioral modulation), the two most critical determinants of stimulation efficacy. Therefore, these findings establish the MNI-protocol as a methodologically sound, practical, and cost-effective alternative to the gold-standard T1-protocol for experimental applications.

Interestingly, the sole distinction between protocols emerged at the anatomical level: the T1-protocol yielded significantly shorter coil-to-cortex distances than the MNI-protocol (Table 1), confirming its superior spatial accuracy. However, this anatomical advantage did not translate into enhanced behavioral modulation or stronger E-field induction, indicating that “spatial closeness” is a necessary but insufficient proxy for TMS efficacy. Notably, coil-to-cortex distances for both protocols fell well within the effective depth range (approximately 2–3 cm) of standard figure-8 coils[35–37] (Table 1). Our data imply that within this operational window, the observed millimeter-scale difference (≈ 2.94 mm) does not cross the biophysical threshold required to produce functional divergence in behavior (Fig. 4A) or E-fields (Fig. 3). While such distance increments may be critical for clinical efficacy[38], in non-clinical experiments with healthy participants, this degree of spatial deviation appears functionally negligible and unlikely to yield measurable differences in stimulation outcomes. Thus, future protocol selection should prioritize functional metrics over anatomical proximity alone.

### 4.2. Significance and implications

The convergence of behavioral and biophysical findings between protocols offers critical insights for TMS research design. First, while T1-based neuronavigation provides superior spatial fidelity, its logistical and financial burdens limit scalability. The MNI-protocol offers a pragmatic, cost-effective alternative that preserves functional efficacy without subject-specific MRI. This is particularly advantageous for pilot studies and large-scale trials, enabling robust data collection and enhanced statistical power. Second, in clinical settings lacking MRI, the MNI-protocol serves as a superior, anatomically informed alternative to probabilistic scalp heuristics (e.g., Beam F3 approach[39]; 5.5 cm rule[40]). Unlike manual methods, MNI registration leverages landmark and head-surface alignment to approximate individual head geometry, enabling greater precision. Furthermore, it offers superior inter-session reproducibility[26, 41, 42], which is critical for multi-session therapeutic protocols that rely on cumulative dosing[41–43]. Thus, the MNI-protocol represents a scalable, anatomically grounded strategy for standardized neuronavigation in both research and clinical domains. Ultimately, our study provides scientific evidence of this utility, establishing the MNI-protocol not as a mere logistical compromise, but as an empirically supported standard for the field.

### 4.3. The dual-space (MS and NS) framework

A pivotal methodological strength of this study is the dual-space analytical framework, strategically designed to evaluate anatomical and electromagnetic indices within their optimal coordinate systems. For the anatomical analysis, we compared coil-to-cortex distances exclusively in MS. By normalizing individual head geometries to a common template, we disentangled protocol-driven targeting errors from inter-subject variability, ensuring a rigorous, standardized comparison of spatial accuracy.

Conversely, for the electromagnetic analysis, prioritizing biophysical validity is paramount. Since E-field propagation is inextricably linked to subject-specific cortical morphology and tissue properties[31, 44], we executed all simulations in each participant’s NS. This approach preserved the intricate individual gyrification patterns that fundamentally shape field distribution. Therefore, this dual-space strategy reconciles group-level comparability with subject-level physiological fidelity, leveraging the distinct advantages of standardization for anatomy and individualization for biophysics.

### 4.4. Limitations

A limitation of this study is its population-specific anatomical constraint. We employed the Caucasian-derived MNI-ICBM152 template[2, 3, 6] for a Korean cohort. Since Western templates do not fully capture non-Western morphology, including variations in brain shape and volume[45], this template-to-subject mismatch likely contributed to the systematic increase in MNI-protocol’s coil-to-cortex distances. While the standard MNI template yielded functionally valid results, future studies involving non-Western populations could enhance anatomical fidelity by adopting ethnicity-specific templates.

## 5. Conclusion

We provide converging evidence that the standardized MNI-protocol is functionally interchangeable with the individualized T1-protocol. Despite systematic anatomical deviations, the MNI-protocol achieved equivalent behavioral modulation and electromagnetic induction. These findings validate MNI-guidance as a scientifically sound alternative to subject-specific MRI. By removing major logistical barriers, this protocol paves the way for scalable, high-throughput TMS applications across diverse experimental and clinical settings.

## Author Contributions

**Hee-Dong Yoon**: Conceptualization, Data curation, Formal analysis, Investigation, Methodology, Resources, Software, Visualization, Writing–original, Writing–review and editing.

**Hyeon-Ae Jeon**: Conceptualization, Funding acquisition, Project administration, Supervision, Writing–original, Writing–review and editing.

## Acknowledgements

We thank Konstantin Weise for his help with electrical field simulations.

