## Supplementary Material for "Standardized MNI-Guided TMS Yields Functional Similarity to Individualized T1-Guidance: Evidence from Behavioral, Anatomical, and Electromagnetic Levels"

Table of Contents

1. Participants - - - - - - - - - - - - - - - - - - - - - - - - - - - - - - - - - - - - - - - - - - - - - - - - - - - - - - - - - - - - - - - - p2
2. Screening questionnaires - - - - - - - - - - - - - - - - - - - - - - - - - - - - - - - - - - - - - - - - - - - - - - - - - - - - - - - - -p3
3. MRI acquisition parameters - - - - - - - - - - - - - - - - - - - - - - - - - - - - - - - - - - - - - - - - - - - - - - - - - - - - - - -p5
4. Number comparison task - - - - - - - - - - - - - - - - - - - - - - - - - - - - - - - - - - - - - - - - - - - - - - - - - - - - - - - - -p5
5. TMS stimulation parameters - - - - - - - - - - - - - - - - - - - - - - - - - - - - - - - - - - - - - - - - - - - - - - - - - - - - - - p6
6. Targeting protocols - - - - - - - - - - - - - - - - - - - - - - - - - - - - - - - - - - - - - - - - - - - - - - - - - - - - - - - - - - - - p7
7. Coordinate transformation validation - - - - - - - - - - - - - - - - - - - - - - - - - - - - - - - - - - - - - - - - - - - - - - - - - p8
8. Table S1. Quantitative validation of coordinate transformation accuracy via round-trip analysis (SPM12) - - - - p10
9. Table S2. Quantitative validation of coordinate transformation accuracy for extracranial space (SPM12)- - - - -p13
10. Table S3. Cross-software validation of coordinate transformation accuracy for the right IPS (FSL) - - - - - - - - p15
11. Neuronavigation system and entry point definition- - - - - - - - - - - - - - - - - - - - - - - - - - - - - - - - - - - - - - - p16
12. Behavioral data preprocessing and trial selection - - - - - - - - - - - - - - - - - - - - - - - - - - - - - - - - - - - - - - p16
13. Calculation of coil-to-cortex distance - - - - - - - - - - - - - - - - - - - - - - - - - - - - - - - - - - - - - - - - - - - - - - p17
14. E-field simulation setup - - - - - - - - - - - - - - - - - - - - - - - - - - - - - - - - - - - - - - - - - - - - - - - - - - - - - - - - p18
15. Figure S1. Individually reconstructed finite element head models for E-field simulations - - - - - - - - - - - - - - p21
16. Real and virtual TMS - - - - - - - - - - - - - - - - - - - - - - - - - - - - - - - - - - - - - - - - - - - - - - - - - - - - - - - - - - -p22
17. ROI definition for E-field quantification - - - - - - - - - - - - - - - - - - - - - - - - - - - - - - - - - - - - - - - - - - - - - - p23
18. Quantification of E-Field spatial extent- - - - - - - - - - - - - - - - - - - - - - - - - - - - - - - - - - - - - - - - - - - - - - - p24
19. Table S4. Comparative quantification of coil-to-cortex distances (mm): T1-protocol versus MNI-protocol - - - p25
20. Figure S2. Individualized E-field distributions for right IPS stimulation: T1- protocol versus MNI-protocol (Real TMS)- - - - - - - - - - - - - - - - - - - - - - - - - - - - - - - - - - - - - - - - - - - - - - - - - - - - - - - - - - - - - - - - - - p26
21. Table S5. Comparative analysis of induced E-field magnitude at the right IPS: T1-protocol versus MNI-protocol (Real TMS)- - - - - - - - - - - - - - - - - - - - - - - - - - - - - - - - - - - - - - - - - - - - - - - - - - - - - - - - - - - - - - p27
22. Table S6 Methodological validation of the virtual TMS approach: Comparison with real TMS at the right IPS-p28
23. References - - - - - - - - - - - - - - - - - - - - - - - - - - - - - - - - - - - - - - - - - - - - - - - - - - - - - - - - - - - - - - - - p29

### 1. Participants

A total of twenty-two healthy university students (12 females, 10 males; age range: 19–24 years; M = 21.14, SD = 1.55) were enrolled in the experimental protocol. All participants were right-handed, as confirmed by the Edinburgh Handedness Inventory (laterality quotient: M = 86.36, SD = 12.93)[1]. While twenty-four individuals were initially assigned recruitment IDs, Subject IDs 18 and 22 withdrew from the study for personal reasons prior to data collection. Consequently, their data are absent from all analyses. To maintain traceability to the original recruitment logs, the original participant IDs (1–24, excluding 18 and 22) were preserved throughout the dataset and manuscript. To ensure experimental safety and data integrity, all candidates underwent a two-stage screening process. First, potential participants were excluded if they presented any contraindications for MRI or TMS, including but not limited to: a history of seizures or epilepsy, implanted ferromagnetic materials or medical devices (e.g., pacemakers, cochlear implants), pregnancy, or use of medications known to lower the seizure threshold. Second, individuals with a history of neurological or psychiatric disorders were excluded to maintain the homogeneity of the healthy cohort. The comprehensive screening questionnaires used in this study are provided in Supplementary Material 2[2, 3]. The experimental protocol was reviewed and approved by the University Institutional Review Board (IRB) and was conducted in strict accordance with the Declaration of Helsinki. Prior to any experimental procedures, all participants provided written informed consent after a full explanation of the study's nature and potential risks. Participants received a fixed monetary compensation for their time and involvement.

### 2. Screening questionnaires

- **MRI Screening Questionnaire (English translation)**

Please answer the following questions. Mark Yes if any item applies to you and provide details where indicated.

1. Cardiac pacemaker
   ☐ Yes ☐ No
2. Implantable cardioverter–defibrillator (ICD) or leads
   ☐ Yes ☐ No
3. Cerebral aneurysm clips
   ☐ Yes ☐ No
4. Cochlear implant or other otologic implants
   ☐ Yes ☐ No
5. Neurostimulator (implantable nerve stimulator)
   ☐ Yes ☐ No
6. Metallic foreign bodies (e.g., pins, screws from accidents)
   ☐ Yes ☐ No
7. Have you ever worked in a metal workshop?
   ☐ Yes ☐ No
8. Do you have any metallic foreign object in the eye?
   ☐ Yes ☐ No
9. History of heart surgery
   ☐ Yes ☐ No
   If yes, please specify:
10. Cardiac or pulmonary assist devices (e.g., heart–lung assist devices)
    ☐ Yes ☐ No
11. Insulin pump
    ☐ Yes ☐ No
12. Orthopedic hardware (e.g., joint clips, screws, plates, pins)
    ☐ Yes ☐ No
13. History of eye surgery or tattooed eyeliner
    ☐ Yes ☐ No
14. For women: intrauterine device (IUD)
    ☐ Yes ☐ No
15. Dental appliances (braces, dentures, oral prostheses)
    ☐ Yes ☐ No
16. Do you currently have tattoos or medicated transdermal patches on the skin?
    ☐ Yes ☐ No
17. Do you have claustrophobia?
    ☐ Yes ☐ No
18. History of surgery
    ☐ Yes ☐ No
    If yes, please provide the reason and the hospital name:
19. For women: are you currently pregnant?
    ☐ Yes ☐ No
20. For women: are you currently breastfeeding?
    ☐ Yes ☐ No
21. Have you previously undergone MRI?
    ☐ Yes ☐ No
22. If you have undergone MRI, did you receive a contrast agent?
    ☐ Yes ☐ No
23. Are you currently taking any medications?
    ☐ Yes ☐ No
    If yes, please list medication name(s) and reason(s):

- **TMS screening questionnaire (English translation)**

Please answer the following questions regarding safety for TMS studies. Mark Yes if any item applies and provide details where indicated.

1. Do you have any implanted devices such as a cardiac pacemaker, portable insulin or medication pump, neurostimulator, shunt system, or cochlear implant?
   ☐ Yes ☐ No
2. Metallic foreign bodies due to accidents or surgery (e.g., pins, screws, fragments)
   ☐ Yes ☐ No
3. Have you undergone surgery within the past two months?
   ☐ Yes ☐ No

   If yes, what surgery did you receive?
4. Implanted metallic joint prosthesis or orthopedic hardware (clips, screws, plates, pins)
   ☐ Yes ☐ No
5. Do you experience discomfort or irregularity in heart rhythm?
   ☐ Yes ☐ No
6. Are you particularly sensitive to sound or does tinnitus interfere with daily life?
   ☐ Yes ☐ No
7. History of epileptic seizures (epilepsy)
   ☐ Yes ☐ No
8. History of febrile seizures between 6 months and 5 years of age
   ☐ Yes ☐ No
9. Family history of epilepsy
   ☐ Yes ☐ No
10. History of loss of consciousness for unclear reasons
    ☐ Yes ☐ No
11. History of diagnosed neurological or psychiatric disorders
    ☐ Yes ☐ No
12. History of head injury (e.g., concussion)
    ☐ Yes ☐ No
13. History of severe headaches requiring treatment
    ☐ Yes ☐ No
14. Do you frequently experience headaches or migraine?
    ☐ Yes ☐ No
15. Do you have a sleep disorder?
    ☐ Yes ☐ No
16. Do you regularly take any medications?
    ☐ Yes ☐ No

    If yes, please list medication name(s):
17. Do you frequently consume alcohol?
    ☐ Yes ☐ No
18. Do you have any chronic medical conditions (e.g., asthma, hypertension)?
    ☐ Yes ☐ No
19. For women: are you currently pregnant or could you be pregnant?
    ☐ Yes ☐ No

Note: This English version is a faithful translation of the original Korean questionnaires provided by the user.

### 3. MRI acquisition parameters

High-resolution T1-weighted magnetic resonance imaging (MRI) scans (T1 images) were acquired using a 3T Siemens MAGNETOM Skyra scanner (Siemens Healthcare, Erlangen, Germany) equipped with a 64-channel phased-array head coil. The T1 images were collected with an MPRAGE sequence (176 sagittal slices; repetition time (TR) = 2300 ms; echo time (TE) = 3.44 ms; flip angle = 9°; field of view (FOV) = 256 mm; voxel size = 1 × 1 × 1 mm; phase-encoding direction = A/P; GRAPPA acceleration factor = 2).

### 4. Number comparison task

For the behavioral task, participants performed a number comparison task. Stimuli consisted of pairs of single-digit Arabic numerals presented bilaterally on a gray background, subtending a horizontal visual angle of approximately 4°. Participants were instructed to identify the numerically larger digit as quickly and accurately as possible while ignoring physical size. Responses were registered via the left or right shift key using the corresponding index finger, spatially mapped to the digit's position on the screen.

Trials were categorized into three conditions based on the congruency between numerical magnitude and physical size:

- Congruent: Numerical and physical dimensions were matched (e.g., the numerically larger digit was also physically larger).
- Incongruent: Numerical and physical dimensions were mismatched (e.g., the numerically larger digit was physically smaller), creating conflict between dimensions.
- Neutral: Digits differed in numerical value but were presented in identical physical sizes.

This factorial design allowed for the quantification of the size congruity effect (SCE), a robust index of cognitive interference arising from cross-dimensional conflict[4-6]. Response times (RTs) were measured from stimulus onset to button press. The SCE was quantified as the difference in RT between incongruent and congruent trials ($RT_{incongruent} - RT_{congruent}$), with larger SCE values indicating greater susceptibility to interference.

The experiment followed a within-subject design spanning three TMS sessions (T1, MNI, and sham). Each session comprised three runs, with 72 trials per run (24 trials per condition: congruent, neutral, incongruent), yielding a total of 216 trials per session and 648 trials per participant. To prevent response strategies, no more than five trials of the same condition were presented consecutively. The inter-trial interval (ITI) was fixed at 6000 ms to accommodate the repetitive TMS (rTMS) protocol. Prior to data collection, participants completed 10 practice trials with feedback. Participants were seated 100 cm from the monitor in a well-lit room, and the total duration of each session was approximately 50 minutes. Stimulus presentation and data logging were controlled using MATLAB R2020a (MathWorks Inc., Natick, MA, USA)[7].

### 5. TMS Stimulation parameters

#### 5.1 Session timing and pulse protocol

To minimize carry-over effects and task-specific learning, the three TMS sessions were separated by a washout period of at least one week (mean inter-session interval = 7.80 days, SD = 3.37). Stimulation was delivered using a MagPro X100 stimulator equipped with a passively cooled MCF-B65 figure-of-eight coil (MagVenture, Farum, Denmark). For the active conditions (T1- and MNI-protocols), we administered event-related repeated TMS (rTMS) consisting of three biphasic pulses delivered at 10 Hz. The pulses were timed at 220, 320, and 420 post-stimulus onset. These latencies were selected to temporally align with the critical time window of parietal event-related potentials (ERPs), which typically onset around 220 ms and reach peak amplitude between 350 and 400 ms during numerical processing[6].

#### 5.2 Stimulation intensity and threshold determination

Stimulation intensity was set at 80% of the individual resting motor threshold (RMT), corresponding to 40–68% of the maximum stimulator output (mean = 49.77%, SD = 8.66; maximum output = 156 A/μs)[8-11]. RMT was defined as the minimum intensity required to elicit visible muscle twitches in the right hand in at least five out of ten consecutive single-pulse stimulations. To locate the motor hotspot, we targeted the left primary motor cortex (M1) using standard MNI coordinates (x = -37, y = -21, z = 58) reported in a prior meta-analysis[12], which were inverse-transformed into each participant's native space (NS) to ensure anatomical precision. Crucially, during both RMT acquisition and experimental stimulation, the coil was positioned perpendicular to the target gyrus (i.e., left M1 and right intraparietal sulcus [IPS]) to maximize the induced electric-field (E-field) strength[13, 14].

5.3 Sham control configuration

The sham session targeted the vertex, a site functionally irrelevant to the task, to control for nonspecific sensory effects. The vertex was manually identified for each participant as the intersection of two midlines: the midpoint between the nasion and inion, and the midpoint between the left and right tragus[15]. To maintain auditory and somatosensory fidelity without inducing significant cortical activation, the coil was flipped 180° away from the scalp[16, 17]. This inversion attenuates the magnetic field intensity by approximately 60%[17], ensuring that the effective stimulation delivered to the cortex remains substantially below the physiological threshold (RMT). In addition, targeting the vertex has been empirically validated as an inert control that exerts no measurable influence on cognitive performance or neural processing during task performance[18, 19].

### 6. Targeting protocols

6.1 T1-protocol (individualized targeting)

The T1-protocol utilized each participant's individual T1-weighted image as the reference for neuronavigation. Co-registration between the physical head and the T1 image was established by aligning anatomical fiducials (nasion, left/right tragus) and additional scalp surface points. Crucially, because the functional target (FT; right IPS) was defined in standard MNI space (MS), we applied an inverse normalization procedure to map the coordinates into each participant's native space (NS). This was achieved using the inverse deformation field derived from the segmentation process in SPM12 (Wellcome Trust Centre for Neuroimaging, University College London, UK; http://www.fil.ion.ucl.ac.uk/spm/)[20], ensuring that the standard MNI coordinate was accurately projected onto the corresponding anatomical locus within the individual brain.

6.2 MNI-protocol (standardized targeting)

In contrast, the MNI-protocol employed the standard MNI-ICBM 152 template for guidance, bypassing the need for individual MRI. In this workflow, the target coordinates were applied directly in MS without cross-space transformation. To map the population-averaged template to the individual participant's head geometry, the neuronavigation software performed a template-to-subject registration. This process utilized the measured scalp landmarks and surface points to estimate a transformation matrix, combining linear (affine) spatial transformations with surface-based deformations (template warping) to approximate the participant's head shape[21].

**7. Coordinate transformation validation**

Coordinate transformation, which is defined as the computational process of converting spatial coordinates from a standardized reference system (e.g., MNI space, MS) to an individual's anatomical space (e.g., native space, NS), and vice versa, was a fundamental prerequisite for the dual-space analytical framework of this study. This bidirectional conversion enabled us to seamlessly integrate standardized targeting comparisons with individualized biophysical modeling. However, precise transformation is critical; even minor registration errors pose a risk of propagating into downstream analyses, potentially confounding the accuracy of coil-to-cortex distance computations and E-field simulations. To rigorously address these concerns and ensure the spatial reliability of our pipeline, we conducted a comprehensive two-stage validation.

First, we evaluated the internal consistency of our primary tool, SPM12, via a "round-trip" transformation procedure. For every participant and across all three cortical targets (right IPS, left DLPFC, and left M1), coordinates were first transformed from MS to NS using inverse deformation fields and subsequently transformed back to MNI space using forward deformation fields. By calculating the

Euclidean displacement between the original and recovered coordinates, we confirmed that transformation errors were negligible for all intracranial targets (Supplementary Material 8). Crucially, because TMS coil positioning involves coordinates located outside the physical head boundary, we extended this validation to arbitrary extracranial points resembling coil locations. This step confirmed that the deformation fields maintained high spatial precision even for extracranial space, ensuring accurate transformation of coil coordinates (Supplementary Material 9). Second, to verify that our results were not driven by algorithm-specific biases inherent to a single software package, we performed a cross-software validation using FMRIB's Software Library (FSL; Center for Functional Magnetic Resonance Imaging of the Brain, University of Oxford, UK)[22] for the right IPS target. While FSL yielded slightly larger variances compared to SPM12, the discrepancies remained consistently within the sub-millimeter range (< 1 mm; Supplementary Material 10). Collectively, these convergent findings, derived from both internal round-trip checks and independent cross-validation, demonstrate that our coordinate transformation pipeline introduced no meaningful spatial shifts, thereby confirming the high fidelity of the spatial data used in our comparative analyses.

**8. Table S1. Quantitative validation of coordinate transformation accuracy via round-trip analysis (SPM12)**

| Subject | Region | MNI_x | MNI_y | MNI_z | MNI_re_x | MNI_re_y | MNI_re_z | Diff_x | Diff_y | Diff_z |
|---|---|---|---|---|---|---|---|---|---|---|
| 1 | Left M1 | -37 | -21 | 58 | -37 | -21 | 58 | 0 | 0 | 0 |
| 1 | Right IPS | 25.63 | -66.73 | 45.28 | 25.63 | -66.73 | 45.28 | 0 | 0 | 0 |
| 1 | Left DLPFC | -30 | 43 | 23 | -30 | 43 | 23 | 0 | 0 | 0 |
| 2 | Left M1 | -37 | -21 | 58 | -37.01 | -21 | 58 | 0.01 | 0 | 0 |
| 2 | Right IPS | 25.63 | -66.73 | 45.28 | 25.63 | -66.73 | 45.27 | 0 | 0 | 0.01 |
| 2 | Left DLPFC | -30 | 43 | 23 | -30 | 43 | 23 | 0 | 0 | 0 |
| 3 | Left M1 | -37 | -21 | 58 | -37 | -21.01 | 58 | 0 | 0.01 | 0 |
| 3 | Right IPS | 25.63 | -66.73 | 45.28 | 25.63 | -66.73 | 45.29 | 0 | 0 | -0.01 |
| 3 | Left DLPFC | -30 | 43 | 23 | -30 | 43 | 23 | 0 | 0 | 0 |
| 4 | Left M1 | -37 | -21 | 58 | -37.01 | -21 | 57.99 | 0.01 | 0 | 0.01 |
| 4 | Right IPS | 25.63 | -66.73 | 45.28 | 25.63 | -66.73 | 45.28 | 0 | 0 | 0 |
| 4 | Left DLPFC | -30 | 43 | 23 | -30 | 43 | 23 | 0 | 0 | 0 |
| 5 | Left M1 | -37 | -21 | 58 | -37 | -21 | 58 | 0 | 0 | 0 |
| 5 | Right IPS | 25.63 | -66.73 | 45.28 | 25.63 | -66.73 | 45.27 | 0 | 0 | 0.01 |
| 5 | Left DLPFC | -30 | 43 | 23 | -30.01 | 43 | 23 | 0.01 | 0 | 0 |
| 6 | Left M1 | -37 | -21 | 58 | -37 | -21 | 58 | 0 | 0 | 0 |
| 6 | Right IPS | 25.63 | -66.73 | 45.28 | 25.63 | -66.73 | 45.27 | 0 | 0 | 0.01 |
| 6 | Left DLPFC | -30 | 43 | 23 | -30 | 43 | 23 | 0 | 0 | 0 |
| 7 | Left M1 | -37 | -21 | 58 | -37 | -21.01 | 58 | 0 | 0.01 | 0 |
| 7 | Right IPS | 25.63 | -66.73 | 45.28 | 25.63 | -66.73 | 45.28 | 0 | 0 | 0 |
| 7 | Left DLPFC | -30 | 43 | 23 | -30 | 43 | 23 | 0 | 0 | 0 |
| 8 | Left M1 | -37 | -21 | 58 | -37 | -21 | 58.01 | 0 | 0 | -0.01 |
| 8 | Right IPS | 25.63 | -66.73 | 45.28 | 25.63 | -66.73 | 45.29 | 0 | 0 | -0.01 |
| 8 | Left DLPFC | -30 | 43 | 23 | -30 | 43 | 23 | 0 | 0 | 0 |
| 9 | Left M1 | -37 | -21 | 58 | -37 | -21.01 | 58 | 0 | 0.01 | 0 |
| 9 | Right IPS | 25.63 | -66.73 | 45.28 | 25.63 | -66.73 | 45.28 | 0 | 0 | 0 |
| 9 | Left DLPFC | -30 | 43 | 23 | -30 | 43 | 23 | 0 | 0 | 0 |
| 10 | Left M1 | -37 | -21 | 58 | -37 | -21 | 58 | 0 | 0 | 0 |
| 10 | Right IPS | 25.63 | -66.73 | 45.28 | 25.63 | -66.73 | 45.28 | 0 | 0 | 0 |
| 10 | Left DLPFC | -30 | 43 | 23 | -30 | 43 | 23 | 0 | 0 | 0 |
| 11 | Left M1 | -37 | -21 | 58 | -37 | -21 | 57.99 | 0 | 0 | 0.01 |
| 11 | Right IPS | 25.63 | -66.73 | 45.28 | 25.63 | -66.74 | 45.28 | 0 | 0.01 | 0 |
| 11 | Left DLPFC | -30 | 43 | 23 | -30 | 43 | 23 | 0 | 0 | 0 |
| 12 | Left M1 | -37 | -21 | 58 | -37 | -20.99 | 58 | 0 | -0.01 | 0 |
| 12 | Right IPS | 25.63 | -66.73 | 45.28 | 25.63 | -66.73 | 45.28 | 0 | 0 | 0 |

| | | | | | | | | | | |
|---|---|---|---|---|---|---|---|---|---|---|
| 12 | Left DLPFC | -30 | 43 | 23 | -30 | 43 | 22.99 | 0 | 0 | 0.01 |
| 13 | Left M1 | -37 | -21 | 58 | -37 | -21 | 58 | 0 | 0 | 0 |
| 13 | Right IPS | 25.63 | -66.73 | 45.28 | 25.63 | -66.73 | 45.28 | 0 | 0 | 0 |
| 13 | Left DLPFC | -30 | 43 | 23 | -30 | 43 | 22.99 | 0 | 0 | 0.01 |
| 14 | Left M1 | -37 | -21 | 58 | -37.01 | -21 | 58 | 0.01 | 0 | 0 |
| 14 | Right IPS | 25.63 | -66.73 | 45.28 | 25.63 | -66.73 | 45.28 | 0 | 0 | 0 |
| 14 | Left DLPFC | -30 | 43 | 23 | -30 | 43 | 23 | 0 | 0 | 0 |
| 15 | Left M1 | -37 | -21 | 58 | -37 | -21 | 58 | 0 | 0 | 0 |
| 15 | Right IPS | 25.63 | -66.73 | 45.28 | 25.63 | -66.73 | 45.28 | 0 | 0 | 0 |
| 15 | Left DLPFC | -30 | 43 | 23 | -30 | 43 | 22.99 | 0 | 0 | 0.01 |
| 16 | Left M1 | -37 | -21 | 58 | -37.01 | -21 | 58.01 | 0.01 | 0 | -0.01 |
| 16 | Right IPS | 25.63 | -66.73 | 45.28 | 25.63 | -66.73 | 45.28 | 0 | 0 | 0 |
| 16 | Left DLPFC | -30 | 43 | 23 | -30 | 43 | 23 | 0 | 0 | 0 |
| 17 | Left M1 | -37 | -21 | 58 | -37 | -21 | 58 | 0 | 0 | 0 |
| 17 | Right IPS | 25.63 | -66.73 | 45.28 | 25.63 | -66.73 | 45.28 | 0 | 0 | 0 |
| 17 | Left DLPFC | -30 | 43 | 23 | -30 | 43.01 | 23 | 0 | -0.01 | 0 |
| 19 | Left M1 | -37 | -21 | 58 | -37 | -21 | 58 | 0 | 0 | 0 |
| 19 | Right IPS | 25.63 | -66.73 | 45.28 | 25.63 | -66.73 | 45.27 | 0 | 0 | 0.01 |
| 19 | Left DLPFC | -30 | 43 | 23 | -29.99 | 43 | 23 | -0.01 | 0 | 0 |
| 20 | Left M1 | -37 | -21 | 58 | -37.01 | -21.01 | 57.99 | 0.01 | 0.01 | 0.01 |
| 20 | Right IPS | 25.63 | -66.73 | 45.28 | 25.63 | -66.73 | 45.28 | 0 | 0 | 0 |
| 20 | Left DLPFC | -30 | 43 | 23 | -29.99 | 43.01 | 23 | -0.01 | -0.01 | 0 |
| 21 | Left M1 | -37 | -21 | 58 | -37.01 | -21 | 57.99 | 0.01 | 0 | 0.01 |
| 21 | Right IPS | 25.63 | -66.73 | 45.28 | 25.63 | -66.73 | 45.28 | 0 | 0 | 0 |
| 21 | Left DLPFC | -30 | 43 | 23 | -30 | 43.01 | 23 | 0 | -0.01 | 0 |
| 23 | Left M1 | -37 | -21 | 58 | -37 | -21 | 58 | 0 | 0 | 0 |
| 23 | Right IPS | 25.63 | -66.73 | 45.28 | 25.63 | -66.73 | 45.28 | 0 | 0 | 0 |
| 23 | Left DLPFC | -30 | 43 | 23 | -30 | 43 | 23 | 0 | 0 | 0 |
| 24 | Left M1 | -37 | -21 | 58 | -37.01 | -21.01 | 58 | 0.01 | 0.01 | 0 |
| 24 | Right IPS | 25.63 | -66.73 | 45.28 | 25.63 | -66.73 | 45.28 | 0 | 0 | 0 |
| 24 | Left DLPFC | -30 | 43 | 23 | -30 | 43 | 23 | 0 | 0 | 0 |

This table presents a coordinate-wise comparison between the original target locations defined in MNI space (MNI_x, MNI_y, MNI_z) and their recovered coordinates following a round-trip transformation (MNI_re_x, MNI_re_y, MNI_re_z). The transformation pipeline involved mapping coordinates from MNI space to individual native space using inverse deformation fields, followed by a reverse mapping back to MNI space using forward deformation fields via SPM12.

- Diff (Diff_x, Diff_y, Diff_z): Calculated as MNI_re – MNI, representing the axis-wise residual displacement (i.e., transformation error) introduced by the bidirectional warping process.
- Values: All measurements are reported in millimeters (mm) and rounded to the nearest hundredth.
- Exclusions: Subject IDs 18 and 22 are omitted as these participants withdrew prior to data collection.

- Abbreviations: M1, primary motor cortex; IPS, intraparietal sulcus; DLPFC, dorsolateral prefrontal cortex.

**9. Table S2. Quantitative validation of coordinate transformation accuracy for extracranial space (SPM12)**

| Subject | Region | MNI_x | MNI_y | MNI_z | MNI_re_x | MNI_re_y | MNI_re_z | Diff_x | Diff_y | Diff_z |
|---|---|---|---|---|---|---|---|---|---|---|
| 1 | Extracranial point | -43 | -110 | 50 | -43 | -110 | 49.99 | 0 | 0 | 0.01 |
| 2 | Extracranial point | -43 | -110 | 50 | -43 | -110 | 50 | 0 | 0 | 0 |
| 3 | Extracranial point | -43 | -110 | 50 | -42.99 | -109.99 | 50 | -0.01 | -0.01 | 0 |
| 4 | Extracranial point | -43 | -110 | 50 | -43 | -110 | 50 | 0 | 0 | 0 |
| 5 | Extracranial point | -43 | -110 | 50 | -43 | -110 | 50 | 0 | 0 | 0 |
| 6 | Extracranial point | -43 | -110 | 50 | -43.01 | -110 | 50 | 0.01 | 0 | 0 |
| 7 | Extracranial point | -43 | -110 | 50 | -43 | -110 | 50 | 0 | 0 | 0 |
| 8 | Extracranial point | -43 | -110 | 50 | -43 | -110 | 50 | 0 | 0 | 0 |
| 9 | Extracranial point | -43 | -110 | 50 | -43 | -110 | 49.99 | 0 | 0 | 0.01 |
| 10 | Extracranial point | -43 | -110 | 50 | -43 | -110 | 50 | 0 | 0 | 0 |
| 11 | Extracranial point | -43 | -110 | 50 | -43 | -110 | 50 | 0 | 0 | 0 |
| 12 | Extracranial point | -43 | -110 | 50 | -43 | -110 | 50 | 0 | 0 | 0 |
| 13 | Extracranial point | -43 | -110 | 50 | -43 | -110 | 50 | 0 | 0 | 0 |
| 14 | Extracranial point | -43 | -110 | 50 | -43 | -110 | 50 | 0 | 0 | 0 |
| 15 | Extracranial point | -43 | -110 | 50 | -43 | -110 | 50 | 0 | 0 | 0 |
| 16 | Extracranial point | -43 | -110 | 50 | -43 | -109.99 | 50 | 0 | -0.01 | 0 |
| 17 | Extracranial point | -43 | -110 | 50 | -43 | -110 | 50 | 0 | 0 | 0 |
| 19 | Extracranial point | -43 | -110 | 50 | -43.01 | -110 | 50 | 0.01 | 0 | 0 |
| 20 | Extracranial point | -43 | -110 | 50 | -43 | -110 | 50 | 0 | 0 | 0 |
| 21 | Extracranial point | -43 | -110 | 50 | -43 | -110 | 50 | 0 | 0 | 0 |
| 23 | Extracranial point | -43 | -110 | 50 | -43 | -110 | 50 | 0 | 0 | 0 |
| 24 | Extracranial point | -43 | -110 | 50 | -43 | -110 | 50 | 0 | 0 | 0 |

Note: This table presents a coordinate-wise comparison between the original target locations defined in MNI space (MNI_x, MNI_y, MNI_z) and their recovered coordinates following a round-trip transformation (MNI_re_x, MNI_re_y, MNI_re_z). The transformation pipeline involved mapping coordinates from MNI space to individual native space using inverse deformation fields, followed by a reverse mapping back to MNI space using forward deformation fields via SPM12.

- Diff (Diff_x, Diff_y, Diff_z): Calculated as MNI_re – MNI, representing the axis-wise residual displacement (i.e., transformation error) introduced by the bidirectional warping process.

- Values: All measurements are reported in millimeters (mm) and rounded to the nearest hundredth.
- Exclusions: Subject IDs 18 and 22 are omitted as these participants withdrew prior to data collection.
- Abbreviations: M1, primary motor cortex; IPS, intraparietal sulcus; DLPFC, dorsolateral prefrontal cortex.

**10. Table S3. Cross-software validation of coordinate transformation accuracy for the right IPS (FSL)**

| Subject | Region | MNI_x | MNI_y | MNI_z | MNI_re_x | MNI_re_y | MNI_re_z | Diff_x | Diff_y | Diff_z |
|---|---|---|---|---|---|---|---|---|---|---|
| 1 | Right IPS | 25.63 | -66.73 | 45.28 | 25.63 | -66.71 | 45.28 | 0.00 | -0.02 | 0.00 |
| 2 | Right IPS | 25.63 | -66.73 | 45.28 | 25.63 | -66.72 | 45.27 | 0.00 | -0.01 | 0.01 |
| 3 | Right IPS | 25.63 | -66.73 | 45.28 | 25.62 | -66.74 | 45.28 | 0.01 | 0.01 | 0.00 |
| 4 | Right IPS | 25.63 | -66.73 | 45.28 | 25.63 | -66.73 | 45.28 | 0.00 | 0.00 | 0.00 |
| 5 | Right IPS | 25.63 | -66.73 | 45.28 | 25.63 | -66.72 | 45.28 | 0.00 | -0.01 | 0.00 |
| 6 | Right IPS | 25.63 | -66.73 | 45.28 | 25.62 | -66.75 | 45.32 | 0.01 | 0.02 | -0.04 |
| 7 | Right IPS | 25.63 | -66.73 | 45.28 | 25.63 | -66.73 | 45.29 | 0.00 | 0.00 | -0.01 |
| 8 | Right IPS | 25.63 | -66.73 | 45.28 | 25.63 | -66.73 | 45.29 | 0.00 | 0.00 | -0.01 |
| 9 | Right IPS | 25.63 | -66.73 | 45.28 | 25.63 | -66.72 | 45.27 | 0.00 | -0.01 | 0.01 |
| 10 | Right IPS | 25.63 | -66.73 | 45.28 | 25.64 | -66.72 | 45.28 | -0.01 | -0.01 | 0.00 |
| 11 | Right IPS | 25.63 | -66.73 | 45.28 | 25.62 | -66.73 | 45.28 | 0.01 | 0.00 | 0.00 |
| 12 | Right IPS | 25.63 | -66.73 | 45.28 | 25.65 | -66.73 | 45.28 | -0.02 | 0.00 | 0.00 |
| 13 | Right IPS | 25.63 | -66.73 | 45.28 | 25.64 | -66.73 | 45.27 | -0.01 | 0.00 | 0.01 |
| 14 | Right IPS | 25.63 | -66.73 | 45.28 | 25.63 | -66.72 | 45.24 | 0.00 | -0.01 | 0.04 |
| 15 | Right IPS | 25.63 | -66.73 | 45.28 | 25.64 | -66.72 | 45.28 | -0.01 | -0.01 | 0.00 |
| 16 | Right IPS | 25.63 | -66.73 | 45.28 | 25.63 | -66.72 | 45.28 | 0.00 | -0.01 | 0.00 |
| 17 | Right IPS | 25.63 | -66.73 | 45.28 | 25.62 | -66.74 | 45.29 | 0.01 | 0.01 | -0.01 |
| 19 | Right IPS | 25.63 | -66.73 | 45.28 | 25.63 | -66.71 | 45.27 | 0.00 | -0.02 | 0.01 |
| 20 | Right IPS | 25.63 | -66.73 | 45.28 | 25.57 | -66.73 | 45.29 | 0.06 | 0.00 | -0.01 |
| 21 | Right IPS | 25.63 | -66.73 | 45.28 | 25.66 | -66.72 | 45.27 | -0.03 | -0.01 | 0.01 |
| 23 | Right IPS | 25.63 | -66.73 | 45.28 | 25.65 | -66.74 | 45.27 | -0.02 | 0.01 | 0.01 |
| 24 | Right IPS | 25.63 | -66.73 | 45.28 | 25.64 | -66.73 | 45.26 | -0.01 | 0.00 | 0.02 |

Note: To verify the robustness of our spatial pipeline and rule out software-specific biases, we replicated the round-trip transformation analysis for the right IPS target using FSL (FMRIB's Software Library). The table compares the original MNI coordinates (MNI_x, MNI_y, MNI_z) with the coordinates recovered after a bidirectional transformation cycle (MNI_re_x, MNI_re_y, MNI_re_z).

- Methodology: Affine transformation matrices and nonlinear warp fields were computed using FSL’s FLIRT (FMRIB's Linear Image Registration Tool) and FNIRT (Nonlinear Image Registration Tool)[23, 24].
- Preprocessing: Prior to registration, T1-weighted images were skull-stripped using FSL’s Brain Extraction Tool (BET) for 17 participants, while original images were used for the remaining 5 participants to assess algorithm performance across different input types.
- Diff (Diff_x, Diff_y, Diff_z): Represents the axis-wise transformation error (MNI_re – MNI).
- Values: All measurements are in millimeters (mm) and rounded to the nearest hundredth.
- Exclusions: Subject IDs 18 and 22 are omitted due to withdrawal.
- Abbreviations: IPS, intraparietal sulcus.

### 11. Neuronavigation system and entry point (EP) definition

We utilized a frameless stereotactic neuronavigation system (Localite TMS Navigator, Localite GmbH, Sankt Augustin, Germany) equipped with an infrared camera-based optical tracking system (Polaris Spectra, NDI, Waterloo, Canada). This setup enabled high-precision, real-time visualization of the TMS coil relative to the targeted cortical regions, ensuring accurate and stable coil positioning throughout the experiment.

#### 11.1 Image registration and surface definition

Depending on the protocol, either the participant's individual T1 image (T1-protocol) or the MNI template (MNI-protocol) was co-registered to the physical head using the Localite registration algorithm[21]. To accurately define the scalp boundary for coil placement, we applied an intensity threshold to the input image (T1 or MNI) to segment the head surface from the background. This threshold was manually adjusted and visually verified during the pre-processing stage to ensure that the calculated head surface faithfully matched the actual scalp contour.

#### 11.2 Entry point (EP) calculation

The EP, defines as the the optimal scalp coordinate for coil placement, was automatically computed to minimize the trajectory distance to the functional target (FT). Using the "Calculate Entry" function, the software identified the scalp vertex yielding the shortest Euclidean path to the specified cortical target[21]. The resulting EP and its trajectory vector were visualized on the interface, serving as the fixed reference anchor for coil positioning across all stimulation sessions.

### 12. Behavioral data preprocessing and trial selection

#### 12.1 Data cleaning and outlier removal

Prior to the analysis, trials with incorrect responses or RTs exceeding ±2.5 SDs from the individual mean were excluded, resulting in the removal of approximately 6% of all trials. The size congruity effect (SCE) was calculated for each session as the difference in mean RT between incongruent and congruent conditions

($RT_{incongruent} - RT_{congruent}$). To enhance reliability and mitigate the distortion caused by rare extreme values [25], we applied a strict trimming procedure, excluding SCE data points beyond ±1.5 SDs from the session mean (approximately 12% exclusion). This step successfully reduced between-subject variability without altering central tendencies, consistent with simulation-based recommendations[25]. Specifically, variability decreased substantially across all sessions (SD reduction: T1: 0.0265 → 0.0193 [−27.17%]; MNI: 0.0317 → 0.0247 [−22.08%]; Sham: 0.0250 → 0.0192 [−23.20%]).

12.2 Restriction to early trials

To maximize sensitivity to TMS-induced disruption, we restricted the primary behavioral analysis to the early phase of each session (first 24 trials). Our behavioral design involved extensive task exposure (648 trials per participant), far exceeding typical TMS study protocols[4, 5]. Such high repetition promotes rapid task automatization[26] and learning[27], which can mask transient TMS effects due to ceiling performance or reduced cognitive demand. By focusing on the initial trials, where TMS effects are known to be most pronounced before substantial task familiarization occurs[28], we avoided practice-related attenuation and ensured a more sensitive assessment of protocol efficacy. This yielded a final dataset of 72 trials per participant across the three TMS sessions (T1, MNI, sham).

**13. Calculation of coil-to-cortex distance**

In this study, "distance computation" refers to the quantification of the Euclidean distance between the scalp and the entry point (EP) and the cortical functional target (FT). The FT represents the intended stimulation site, while the EP represents the optimal coil location on the scalp.

To ensure a standardized comparison that controls for individual variability in head size and shape, all final distance computations were performed within the common MNI space (MS). The computation procedures for each protocol were as follows:

- MNI-Protocol: The workflow for the MNI-protocol was straightforward because both the functional target (FT) and the calculated entry point (EP) were already defined within the same

MNI coordinate system (MS). Since the neuronavigation system determined the EP directly based on the MNI template, no spatial transformation was required. We simply calculated the Euclidean distance between the coordinates of the FT and the EP in MS.

- T1-Protocol: Since the T1-protocol operates in the participant's native space (NS), a multi-step transformation pipeline was necessary to allow for comparison in MS:

  (1) Target Individualization (MS → NS): The standardized FT coordinates were first transformed from MS to each participant's NS using inverse normalization (via SPM12 inverse deformation fields).

  (2) EP Definition (in NS): Using the Localite system's planning mode, the EP was identified in NS as the point on the scalp providing the shortest trajectory to the individualized FT.

  (3) EP Standardization (NS → MS): To compare this EP against the MNI-protocol, the resulting native EP coordinates were transformed back into MS using forward normalization (via SPM12 forward deformation fields).

Finally, for both protocols, the Euclidean distances between the EP and FT in MS was computed using the following formula[29]:

$$Distance_{point1-point2} = \sqrt{(x_{point1} - x_{point2})^2 + (y_{point1} - y_{point2})^2 + (z_{point1} - z_{point2})^2}$$

**14. E-field simulation setup**

14.1 Implementation and technical parameters

For E-field simulation, we used SimNIBS v3.2.1[30, 31], an open-source software platform designed for the realistic modeling of non-invasive brain stimulation. SimNIBS employs a finite element method (FEM) to compute high-resolution E-field distributions based on individual anatomical MRI data. To implement this modeling approach, subject-specific head models were constructed from T1-weighted MRI scans using the headreco pipeline in SimNIBS[32, 33]. Each resulting model consisted of approximately 1.7 million nodes and 9.9 million tetrahedra. Tissue compartments were segmented into five distinct types with the following default conductivity values: scalp (0.465 S/m), skull (0.010 S/m), gray matter (0.276 S/m), white matter (0.126 S/m), and cerebrospinal fluid (1.654 S/m)[13, 32, 34]. This detailed segmentation enabled the anatomically precise estimation of electromagnetic propagation through heterogeneous tissues.

To standardize stimulation strength across individuals, the experimental TMS intensity (% of maximum stimulator output) was converted to the time derivative of the coil current (*dI/dt*, in A/μs), which represents the temporal rate of change in magnetic flux[35]. Simulations were initially performed at a unit intensity (1 A/μs) and subsequently linearly scaled to reflect each participant's specific *dI/dt* values.

14.2 Conceptual framework and biophysical validity

Central to our simulation framework is the prioritization of high-fidelity, individualized modeling, deliberately designed to control for the substantial physiological variability that often confounds TMS research. Even when targeting the identical coordinates in MS, the physiological impact of TMS varies profoundly across individuals due to differences in cortical morphology and brain geometry[36, 37]. Consequently, accurate E-field estimation necessitates subject-specific head models that account for tissue properties and unique gyrification patterns, which fundamentally dictate field propagation[14, 33, 38]. Beyond anatomical modeling, we placed equal emphasis on participant-tailored coil optimization[39]. Mere positional accuracy at the scalp is insufficient; inaccuracies in coil orientation can lead to off-target stimulation, compromising both interpretability and reproducibility[40, 41]. To mitigate this risk, we enforced a consistent, anatomically guided strategy for all simulations (both real and virtual): aligning the coil perpendicular to the local cortical fold[13, 14]. This physiologically grounded approach ensured that E-field induction was optimized for each participant's specific anatomy, providing a highly controlled and valid basis for evaluating protocol-level differences.

14.3 Spatial coordination and ROI analysis

All E-field simulations were executed in the participant's native space (NS). Consequently, coil poses (position and orientation) for both protocols were specified within the NS coordinate system. For the T1-

protocol, the entry point (EP-T1p) was naturally defined in NS. In contrast, for the MNI-protocol, the EP-MNIp, initially calculated in MNI space (MS), was transformed into NS (EP-MNIp in NS) to replicate the physical stimulation geometry on the individual head model. Coils were positioned on the scalp based on these EPs, and poses were defined accordingly (physical coil poses for real TMS and manually constructed poses for virtual TMS; see Supplementary Material 16 for detailed descriptions).

For quantitative analysis, we defined a region of interest (ROI) for each target (the right IPS, left DLPFC, and left M1) as a 5-mm radius sphere[32] centered on the function target (FT) coordinates in NS (transformed from the predefined FT in MS). To ensure anatomical specificity, ROI sampling was strictly constrained to gray matter elements within the tetrahedral mesh. For each ROI, the mean E-field magnitude was calculated and compared across protocols. Finally, for group-level visualization, the individually modeled E-field maps were resampled to the FreeSurfer template brain (fsaverage) and averaged across participants[42].

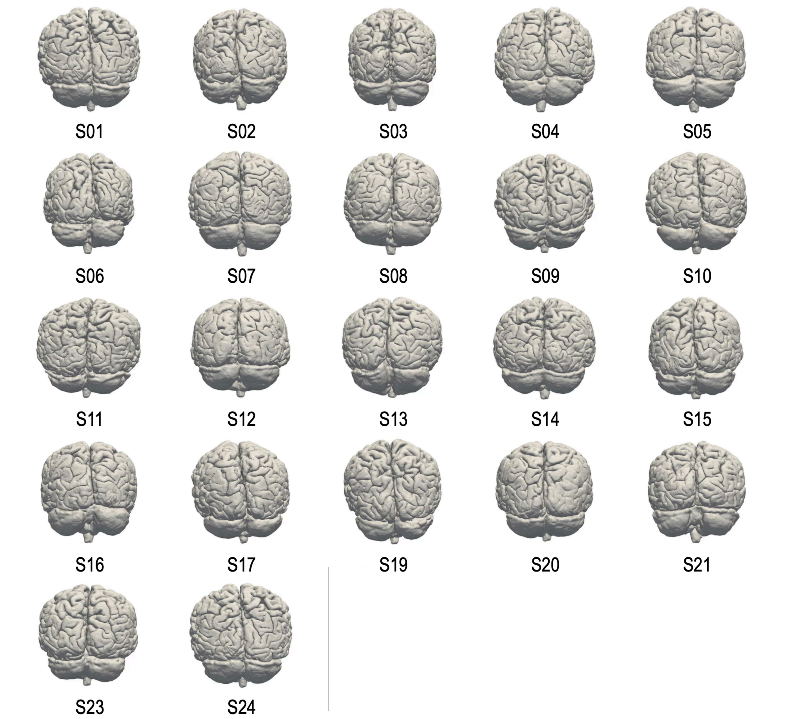


**15. Figure S1. Individually reonstructed finite element head models for E-field simulations.** Subject-specific head models were constructed from high-resolution T1-weighted MRI scans using the headreco pipeline in SimNIBS (v.3.2.1). Each panel displays the posterior view of the reconstructed cortical surface mesh for individual participants, illustrating the distinct inter-subject variability in gyral morphology preserved in the Native Space simulations. Note: Subject IDs 18 and 22 are omitted due to participant withdrawal prior to data acquisition.

**16. Real and virtual TMS**

To enable comprehensive E-field comparisons across multiple cortical targets (right intraparietal sulcus [IPS], left dorsolateral prefrontal cortex [DLPFC], and left primary motor cortex [M1]) despite the limitation that real-world physical coil poses were empirically recorded only for the right IPS during the behavioral experiment, we implemented a dual-simulation approach: Real TMS and Virtual TMS.

16.1 Real TMS (empirically driven simulation)

This approach reconstructed the exact stimulation conditions delivered during the experiment. During the behavioral sessions, the physical TMS coil was positioned on the scalp according to the protocol-specific entry point (EP) coordinates in native space (EP-T1p in NS or EP-MNIp in NS). The precise coil position and orientation were tracked and recorded by the neuronavigation system. Consistent with standard practice, the coil was oriented perpendicular to the local gyrus to maximize the induced E-field strength [13, 14]. These empirically recorded coil poses were subsequently imported into the subject-specific head models to compute the E-field. In this study, real TMS simulations were restricted to the right IPS because this was the only region where physical stimulation occurred and where real-world coil pose tracking was available.

16.2 Virtual TMS (manually defined simulation)

To extend our analysis to regions without empirical tracking data (left DLPFC and left M1) and to ensure methodological consistency across all targets, we employed a "Virtual TMS" approach. Here, coil poses were manually defined on subject-specific head models using the SimNIBS graphical interface. Virtual coil placement was strictly anchored to the protocol-specific EP coordinates in NS. The virtual coil was positioned tangent to the scalp at the EP and oriented perpendicular to the underlying local gyrus, replicating the orientation strategy used in the real experiment. This virtual approach was applied to all three regions (right IPS, left DLPFC and left M1) to facilitate a uniform cross-regional comparison.

16.3 Validation of the virtual approach

Prior to adopting virtual TMS as the primary method for our multi-regional analysis, we validated its accuracy at the right IPS by comparing E-field estimates derived from virtual TMS (manual definition) against those from real TMS (empirical recording). This validation was performed specifically using the T1-protocol, which served as the most stringent reference standard because if utilizes the participant's own anatomy for both targeting and head modeling within a unified native space (NS). This design minimized potential confounds related to template-to-subject registration, allowing us to isolate the impact of coil pose specification methods. This analysis confirmed that virtual TMS successfully reproduced the E-field magnitude and distribution profiles obtained from real TMS at the right IPS (see Fig. 5 and Supplementary Material 22). This empirical validation justified the use of the virtual TMS approach for assessing protocol interchangeability across the broader cortical regions.

16.4 Simulation matrix

In total, eight simulations were performed for each participant:

- Real TMS (2 simulations): Right IPS under T1- and MNI-protocols (used for protocol comparison at the behavioral target).
- Virtual TMS (6 simulations): Right IPS, left DLPFC, and left M1 under both T1- and MNI-protocols (used for evaluating regional generalizability).

**17. ROI definition for E-field quantification**

A critical methodological consideration was how to define the target region for quantifying electromagnetic outcomes. While previous studies have employed anatomically defined regions of interest (ROIs) derived from probabilistic atlases[15], such as the SPM Anatomy Toolbox[43], this approach is inherently confounded by inter-individual variability in cortical anatomy, including differences in the size, shape, and precise location of functional areas. Such variability can introduce bias when comparing E-field magnitudes across subjects.

To eliminate this confound and enable a strictly standardized comparison, we operationalized the ROI as a fixed 5-mm radius sphere centered on each subject-specific functional target (FT). This geometric standardization ensures that E-field magnitude is sampled within a tightly localized and identical volumetric space across all participants and protocols. The 5-mm radius was selected as an optimal parameter to balance focality with signal stability, providing a robust average of the E-field while adhering to established precedents in E-field modeling[44, 45].

**18. Quantification of E-Field spatial extent**

Complementing the analysis of E-field magnitude, we quantified the spatial extent of the induced field to assess potential differences in stimulation focality. To define the area of peak induction, we generated a threshold mask on the cortical surface based on the group-averaged E-field maps. Following the methodology established by Opitz et al.[14], this mask encompassed all triangular surface elements of the tetrahedral mesh where the E-field intensity exceeded 50% of the group-maximum magnitude. The spatial extent was then computed as the total surface area (in $mm^2$) of these suprathreshold elements. While the 50% cutoff represents a fixed methodological criterion, it provided a consistent normalization standard, enabling a robust relative comparison of the high-intensity field spread between the T1- and MNI-protocols.

**19. Table S4. Comparative quantification of coil-to-cortex distances (mm): T1-protocol versus MNI-protocol.**

| | Right IPS | | Left DLPFC | | Left M1 | |
|---|---|---|---|---|---|---|
| **Subject** | **T1 distance** | **MNI distance** | **T1 distance** | **MNI distance** | **T1 distance** | **MNI distance** |
| 1 | 28.72 | 33.87 | 23.60 | 27.83 | 27.02 | 28.80 |
| 2 | 29.71 | 32.79 | 24.25 | 27.55 | 26.12 | 28.85 |
| 3 | 28.64 | 32.54 | 23.26 | 27.69 | 25.28 | 32.04 |
| 4 | 30.56 | 33.78 | 24.48 | 28.48 | 24.61 | 28.82 |
| 5 | 30.53 | 33.19 | 32.65 | 28.33 | 33.25 | 28.86 |
| 6 | 30.65 | 32.78 | 22.19 | 27.68 | 27.02 | 29.18 |
| 7 | 29.36 | 33.58 | 24.33 | 27.96 | 27.40 | 29.20 |
| 8 | 30.93 | 32.66 | 24.48 | 28.54 | 26.18 | 28.84 |
| 9 | 29.24 | 32.52 | 24.54 | 27.47 | 27.56 | 28.82 |
| 10 | 30.65 | 32.92 | 25.73 | 27.56 | 25.67 | 32.08 |
| 11 | 30.75 | 32.86 | 25.98 | 28.17 | 27.61 | 28.91 |
| 12 | 30.86 | 32.83 | 24.23 | 27.71 | 25.43 | 28.79 |
| 13 | 31.51 | 32.82 | 24.97 | 27.80 | 26.44 | 28.89 |
| 14 | 31.16 | 32.66 | 24.73 | 27.57 | 26.30 | 28.92 |
| 15 | 30.82 | 32.89 | 23.66 | 28.11 | 26.53 | 29.13 |
| 16 | 29.8 | 32.94 | 24.57 | 27.57 | 24.77 | 28.90 |
| 17 | 18.89 | 32.68 | 26.65 | 27.75 | 32.58 | 28.75 |
| 19 | 30.67 | 32.94 | 23.85 | 27.53 | 25.78 | 28.80 |
| 20 | 28.59 | 32.63 | 24.89 | 27.95 | 25.90 | 29.07 |
| 21 | 29.48 | 32.77 | 24.31 | 27.73 | 24.87 | 28.82 |
| 23 | 29.57 | 32.82 | 24.50 | 27.60 | 26.33 | 28.85 |
| 24 | 29.6 | 32.86 | 23.91 | 27.85 | 25.58 | 28.82 |
| **Mean distance** | 29.58 | 32.92 | 24.81 | 27.84 | 26.74 | 29.19 |
| **Difference between mean distances** | 3.35 | | 3.03 | | 2.45 | |

Note: This table details the Euclidean distance between the Functional Target (FT) and the protocol-specific scalp Entry Point (EP), calculated within the standardized MNI Space (MS) to facilitate direct comparison (EP-T1p in MS or EP-MNIp in MS). Data are provided for three cortical regions: right IPS, left DLPFC, and left M1.

- Results: Mean distances and between-protocol differences are summarized at the bottom. Consistent with the main analysis, the MNI-protocol yielded systematically larger distances compared to the T1-protocol across all regions, indicating that the T1-protocol offers superior spatial proximity to the cortical target.
- Values: All measurements are in millimeters (mm) and rounded to the nearest hundredth.
- Exclusions: Subject IDs 18 and 22 are omitted due to withdrawal.
- Abbreviations: IPS, intraparietal sulcus; DLPFC, dorsolateral prefrontal cortex; M1, primary motor cortex; FT, functional target; EP, entry point; MS, MNI space.

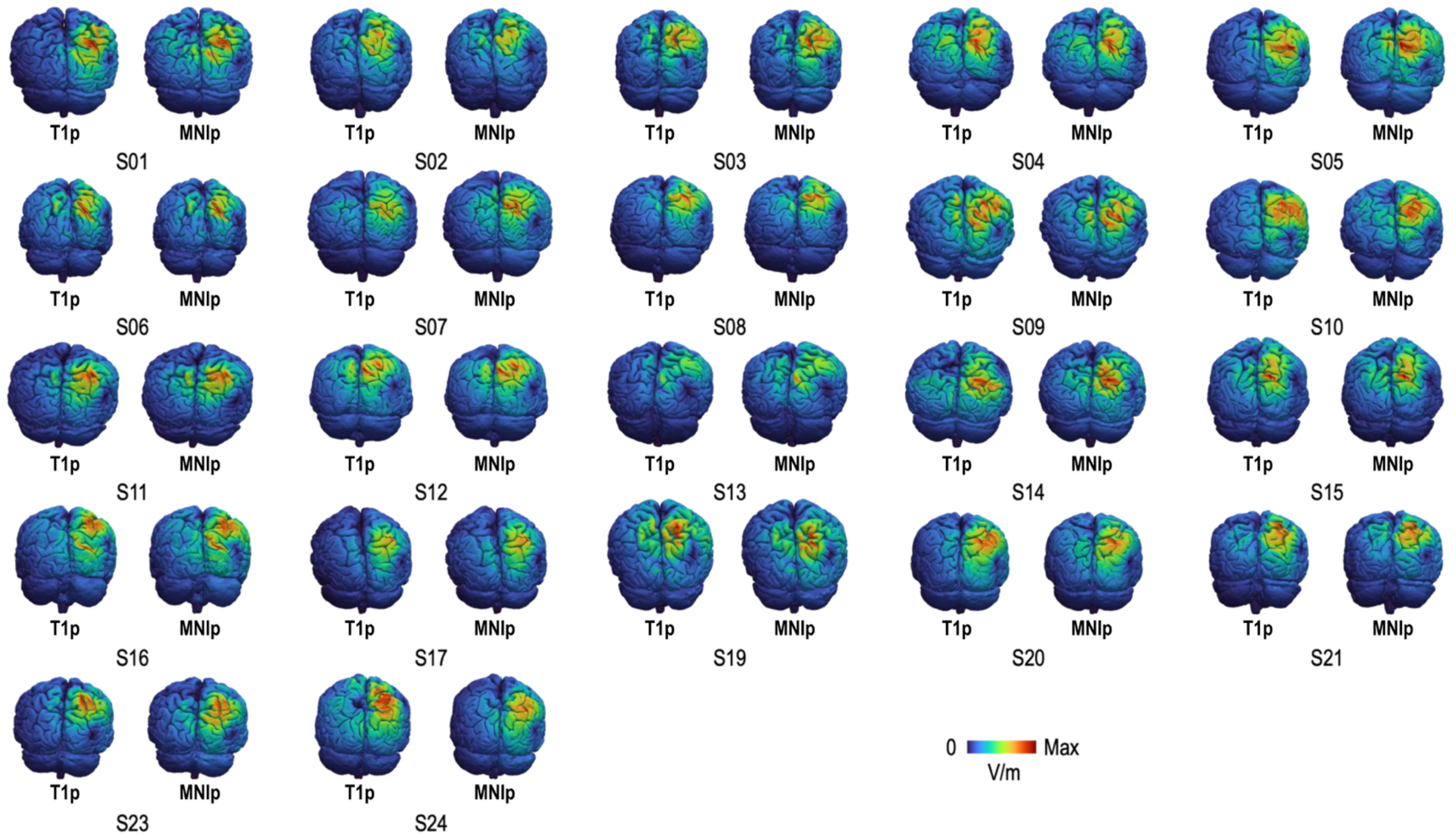


**20. Figure S2. Individualized E-field distributions for right IPS stimulation: T1- protocol versus MNI-protocol (Real TMS).** This figure displays the subject-specific E-field magnitude maps simulated using SimNIBS (v3.2.1) for the right intraparietal sulcus (IPS) target. Simulations were conducted under the "Real TMS" framework, incorporating the actual coil poses (position and orientation) empirically recorded during the behavioral experiment.

- Layout: Each pair represents a single participant, showing the posterior view of the cortical surface for the T1-protocol (T1p, left) and the MNI-protocol (MNIp, right).
- Color Scaling: The colormap indicates the induced E-field magnitude (V/m). To visualize the spatial distribution pattern clearly for each individual, the scale is normalized from 0 V/m to each participant's specific maximum E-field intensity.

Note: Subject IDs 18 and 22 are omitted due to withdrawal prior to data collection.

**21. Table S5. Comparative analysis of induced E-field magnitud at the right IPS: T1-protocol versus MNI-protocol (Real TMS)**

| Subject | Real T1 | Real MNI |
|---|---|---|
| 1 | 39.65 | 34.18 |
| 2 | 19.61 | 20.73 |
| 3 | 31.50 | 29.83 |
| 4 | 25.45 | 24.65 |
| 5 | 29.26 | 30.89 |
| 6 | 28.24 | 26.04 |
| 7 | 29.16 | 27.59 |
| 8 | 27.24 | 25.02 |
| 9 | 19.03 | 18.50 |
| 10 | 32.28 | 36.32 |
| 11 | 35.04 | 43.96 |
| 12 | 18.63 | 21.03 |
| 13 | 17.36 | 16.94 |
| 14 | 25.52 | 33.99 |
| 15 | 22.20 | 28.59 |
| 16 | 27.56 | 23.99 |
| 17 | 58.96 | 38.05 |
| 19 | 28.20 | 28.28 |
| 20 | 25.82 | 27.59 |
| 21 | 24.37 | 29.93 |
| 23 | 33.39 | 34.03 |
| 24 | 40.38 | 35.92 |
| **Mean E-Field (SD)** | 29.04 (9.14) | 28.91 (6.73) |

Note: This table presents the participant-wise mean E-field magnitude (V/m) simulated at the right intraparietal sulcus (IPS). Simulations were conducted under the real TMS framework, utilizing coil poses (position and orientation) empirically recorded during the behavioral sessions.

- Quantification: E-field values were extracted from a 5-mm radius spherical ROI, centered on the functional target in native space (NS) and strictly constrained to gray matter elements.
- Results: Mean E-field magnitudes and the across-participant standard deviations (SD) are summarized at the bottom. A paired-sample t-test confirmed no statistically significant difference in induced field strength between the T1- and MNI-protocols, indicating biophysical equivalence at the primary target.
- Values: All measurements are in V/m and rounded to the nearest hundredth.
- Exclusions: Subject IDs 18 and 22 are omitted due to withdrawal.
- Abbreviations: IPS, intraparietal sulcus.

**22. Table S6. Methodological validation of the virtual TMS approach: Comparison with real TMS at the right IPS.**

| Subject | Real TMS | Virtual TMS |
|---|---|---|
| 1 | 39.65 | 43.45 |
| 2 | 19.61 | 24.02 |
| 3 | 31.50 | 30.77 |
| 4 | 25.45 | 26.10 |
| 5 | 29.26 | 25.16 |
| 6 | 28.24 | 32.00 |
| 7 | 29.16 | 28.17 |
| 8 | 27.24 | 28.82 |
| 9 | 19.03 | 20.59 |
| 10 | 32.28 | 35.80 |
| 11 | 35.04 | 36.09 |
| 12 | 18.63 | 21.52 |
| 13 | 17.36 | 22.81 |
| 14 | 25.52 | 33.06 |
| 15 | 22.20 | 21.91 |
| 16 | 27.56 | 28.46 |
| 17 | 58.96 | 38.12 |
| 19 | 28.20 | 30.11 |
| 20 | 25.82 | 32.04 |
| 21 | 24.37 | 27.22 |
| 23 | 33.39 | 34.83 |
| 24 | 40.38 | 42.80 |
| **Mean E-Field (SD)** | **29.04 (9.14)** | **30.18 (6.51)** |

Note: To justify the use of manually defined coil poses for regions lacking empirical tracking data, we compared E-field magnitudes (V/m) simulated from real TMS (empirically recorded coil poses) against virtual TMS (manually configured coil poses) at the right IPS using the T1-protocol.

- Definitions: Real TMS values represent the actual stimulation delivered during the experiment, while virtual TMS values represent the idealized targeting configuration modeled on subject-specific anatomy.
- Results: Comparison of the mean E-field magnitudes reveals high concordance between the two approaches, confirming that the virtual TMS framework serves as a reliable proxy for empirical stimulation.
- Values: All measurements are in V/m and rounded to the nearest hundredth.
- Exclusions: Subject IDs 18 and 22 are omitted due to withdrawal.
- Abbreviations: IPS, intraparietal sulcus.

[23] Andersson JL, Jenkinson M, Smith S. Non-linear registration, aka Spatial normalisation FMRIB technical

report TR07JA2. FMRIB Analysis Group of the University of Oxford 2007;2(1):e21.
[24] Jenkinson M, Bannister P, Brady M, Smith S. Improved optimization for the robust and accurate linear registration and motion correction of brain images. Neuroimage 2002;17(2):825-41.
[25] Ratcliff R. Methods for dealing with reaction time outliers. Psychological bulletin 1993;114(3):510.
[26] Schneider W, Shiffrin RM. Controlled and automatic human information processing: I. Detection, search, and attention. Psychological review 1977;84(1):1.
[27] Karpicke Jeffrey D, Roediger Henry L. The Critical Importance of Retrieval for Learning. Science 2008;319(5865):966-8.
[28] Jansma JM, van Raalten TR, Boessen R, Neggers SFW, Jacobs RHAH, Kahn RS, et al. fMRI guided rTMS evidence for reduced left prefrontal involvement after task practice. PLoS One 2013;8(12):e80256.
[29] Theodoroff SM, Stevens AA, McMillan G, Pettersson DR, Woodward W, Folmer RL. MRI Verification of a 10–20 Targeting Protocol Used During Transcranial Magnetic Stimulation Sessions for Tinnitus. Brain topography 2018;31(4):690-9.
[30] Saturnino GB, Madsen KH, Thielscher A. Electric field simulations for transcranial brain stimulation using FEM: an efficient implementation and error analysis. Journal of neural engineering 2019;16(6):066032.
[31] Thielscher A, Antunes A, Saturnino GB. Field modeling for transcranial magnetic stimulation: a useful tool to understand the physiological effects of TMS? *2015 37th annual international conference of the IEEE engineering in medicine and biology society (EMBC)*. IEEE; 2015:222-5.
[32] Saturnino GB, Puonti O, Nielsen JD, Antonenko D, Madsen KH, Thielscher A. SimNIBS 2.1: a comprehensive pipeline for individualized electric field modelling for transcranial brain stimulation. Brain and Human Body Modeling 2019:3-25.
[33] Saturnino GB, Madsen KH, Thielscher A. Efficient electric field simulations for transcranial brain stimulation. bioRxiv 2019:541409.
[34] Wagner TA, Zahn M, Grodzinsky AJ, Pascual-Leone A. Three-dimensional head model simulation of transcranial magnetic stimulation. IEEE Transactions on Biomedical Engineering 2004;51(9):1586-98.
[35] Basser PJ, Roth BJ. Stimulation of a myelinated nerve axon by electromagnetic induction. Medical & Biological Engineering & Computing 1991;29(3):261-8.
[36] Syeda F, Magsood H, Lee E, El-Gendy A, Jiles DC, Hadimani R. Effect of anatomical variability in brain on transcranial magnetic stimulation treatment. Aip Advances 2017;7(5):056711.
[37] Wang J, Zhai J, Wang Y, Lin J, Pan D, Li L. Cortical fold geometry modulates transcranial magnetic stimulation electric field strength and peak displacement. Scientific Reports 2025;15(1):19361.
[38] Bungert A, Antunes A, Espenhahn S, Thielscher A. Where does TMS Stimulate the Motor Cortex? Combining Electrophysiological Measurements and Realistic Field Estimates to Reveal the Affected Cortex Position. Cereb Cortex 2017;27(11):5083-94.
[39] Trillenberg P, Bremer S, Oung S, Erdmann C, Schweikard A, Richter L. Variation of stimulation intensity in transcranial magnetic stimulation with depth. Journal of neuroscience methods 2012;211(2):185-90.
[40] Laakso I, Hirata A, Ugawa Y. Effects of coil orientation on the electric field induced by TMS over the hand motor area. Physics in Medicine & Biology 2013;59(1):203.
[41] Gomez LJ, Dannhauer M, Peterchev AV. Fast computational optimization of TMS coil placement for individualized electric field targeting. Neuroimage 2021;228:117696.
[42] Wu J, Ngo GH, Greve D, Li J, He T, Fischl B, et al. Accurate nonlinear mapping between MNI volumetric and FreeSurfer surface coordinate systems. Human brain mapping 2018;39(9):3793-808.
[43] Eickhoff SB, Stephan KE, Mohlberg H, Grefkes C, Fink GR, Amunts K, et al. A new SPM toolbox for combining probabilistic cytoarchitectonic maps and functional imaging data. Neuroimage 2005;25(4):1325-35.
[44] Numssen O, Kuhnke P, Weise K, Hartwigsen G. Electric-field-based dosing for TMS. Imaging Neuroscience 2024;2:1-12.
[45] Zhang BBB, Stöhrmann P, Godbersen GM, Unterholzner J, Kasper S, Kranz GS, et al. Normal component of TMS-induced electric field is correlated with depressive symptom relief in treatment-resistant depression. Brain Stimulation: Basic, Translational, and Clinical Research in Neuromodulation 2022;15(5):1318-20.
